# Spike-phase coupling of subthalamic neurons to posterior opercular cortex predicts speech sound accuracy

**DOI:** 10.1101/2023.10.18.562969

**Authors:** Matteo Vissani, Alan Bush, Witold J. Lipski, Latané Bullock, Petra Fischer, Clemens Neudorfer, Lori L. Holt, Julie A. Fiez, Robert S. Turner, R. Mark Richardson

## Abstract

Speech provides a rich context for understanding how cortical interactions with the basal ganglia contribute to unique human behaviors, but opportunities for direct intracranial recordings across cortical-basal ganglia networks are rare. We recorded electrocorticographic signals in the cortex synchronously with single units in the basal ganglia during awake neurosurgeries where subjects spoke syllable repetitions. We discovered that individual STN neurons have transient (200ms) spike-phase coupling (SPC) events with multiple cortical regions. The spike timing of STN neurons was coordinated with the phase of theta-alpha oscillations in the posterior supramarginal and superior temporal gyrus during speech planning and production. Speech sound errors occurred when this STN-cortical interaction was delayed. Our results suggest that the STN supports mechanisms of speech planning and auditory-sensorimotor integration during speech production that are required to achieve high fidelity of the phonological and articulatory representation of the target phoneme. These findings establish a framework for understanding cortical-basal ganglia interaction in other human behaviors, and additionally indicate that firing-rate based models are insufficient for explaining basal ganglia circuit behavior.

## Introduction

In everyday conversation, humans understand and produce speech with remarkable accuracy and speed. Fluent speech requires the coordination and sequential movement of oral articulators on the order of milliseconds^1,2^. After conceptualization and sequencing in the language network, the brain sends complex motor signals to the larynx, lips, jaw, and tongue, which activate and shape the vocal tract. Brain networks with both cortical and subcortical nodes subserve the coordination of speech. Cognitive neuroscience has delineated the cortical network subserving speech, but less is known about the subcortical contributions, and even less is known about how different nodes in the network transmit and share information.

Despite the impact of speech impairments in basal ganglia disorders such as Parkinson’s Disease (PD) and dystonia, how the basal ganglia participate in speech processing remains unclear. Much of the existing literature has emphasized expansive cortical networks and their dynamics, with abstract references to the basal ganglia^2–4^ because there is little empiric data on which to build models. Indeed, the limitations of functional magnetic resonance imaging (fMRI) for measuring the activity of individual basal ganglia nuclei preclude the explicit integration of subcortical nodes within current theoretical frameworks ^1,5–7^. The scarcity of invasive subcortical data during speech production presents a knowledge gap that hinders our broader understanding of basal ganglia-cortical networks and limits the expansion of current cortico-centric models of speech production (e.g., Directions Into Velocities of Articulators (DIVA) ^5^ and State feedback control (SFC) ^6^).

Recordings from awake Deep Brain Stimulation (DBS) surgeries offer a rare window into human basal ganglia electrophysiology. The subthalamic nucleus (STN) is a common target in DBS for Parkinson’s disease. The STN occupies a privileged role in the basal ganglia thalamo-cortical network, receiving monosynaptic cortical projections through the hyperdirect pathway ^8–10^, in addition to striatal input through the indirect pathway, positioning it as a hub for the temporal integration and spatial compression of information originating from multiple cortical areas ^11,12^. The discovery of single unit and population level activity in the subthalamic nucleus that tracks multiple aspects of speech production ^13–16^ and emerging evidence for anatomical ^9^ and functional connectivity ^17^ between the STN and sensorimotor and auditory cortical areas, raises the question of how STN and cortex interact to mediate speech-related behavior.

In particular, our group established a protocol for high-density electrocorticography (ECoG) to record local field potentials across cortical areas known to participate in speech perception, planning, and production, simultaneous with recording STN single-unit activity during DBS surgery. This paradigm allowed us to study cortico-subcortical spike-phase coupling (SPC) – the relationship between STN spike timing and cortical oscillation phase ^18–21^. We tested the hypothesis that STN neurons are engaged in inter-areal SPC with cortical regions to exercise control on the planning and implementation of the desired articulatory sequence. This unique approach allowed investigation of the spatiotemporal organization of cortico-subcortical interactions with millisecond resolution during speech production. We examined (1) whether SPC was modulated during a syllable triplet repetition task, (2) the spatial and behavioral (temporal) specificity of these relationships and (3) if SPC reflected participants’ ability to utter the desired speech sound.

## Results

We studied intracranial recordings in 24 participants (see Table S1 for demographic details) undergoing STN-DBS surgery for the treatment of Parkinson’s disease. High-density electrocorticography (ECoG) across the left ventral sensorimotor cortex, superior temporal gyrus, and inferior frontal regions were recorded simultaneously with single neuron activity from the STN (Figure 1A). Following the presentation of an auditory cue of a syllable triplet comprised of three unique phototactically-legal consonant-vowel (CV) syllables, participants were instructed to repeat the syllable triplet (speech production) at their own pace into an omnidirectional microphone (64 recording sessions, 2.67 ± 0.62 sessions, 379.88 ± 99.49 trials on average across participants) (Figure 1B). Participants produced the CV-CV-CV sequences in 1.35 ± 0.41 s with a phonetic accuracy of 56.41 ± 26.91% (a triplet was considered inaccurate if any phoneme was off-target). Phonetic errors included consonant substitutions, such as the transformation of plosives into fricatives (e.g., */g/*->*/v/*) and vice-versa (66.26 ± 20.68%), vowel substitution (8.08 ± 11.56 %) and omissions (25.67 ± 19.88 %) (Figure S1). We did not observe an effect of the syllable position in the triplet on phonetic error frequency.

**Figure 1:**
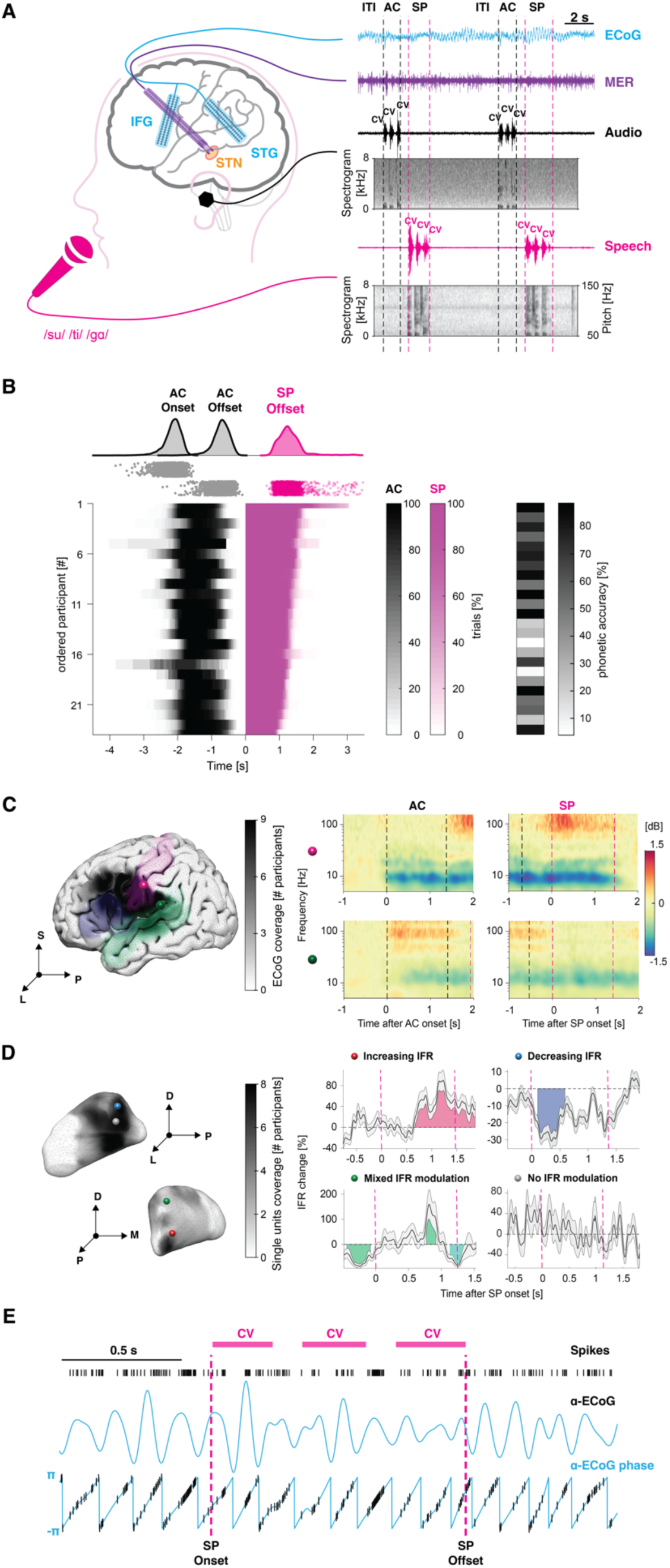
Quantification of STN-cortex spikephase coupling during an intraoperative syllable triplet repetition task. (**A**) Illustration of the syllable triplet repetition task. Participants were instructed to repeat unique consonant-vowel (“CV”) syllable triplets (magenta). The auditory stimuli were presented through earphones (black). High-density electrocorticography (ECoG) strips were placed in auditory and sensorimotor areas through the burr hole (cyan). Microelectrode recordings were acquired in the subthalamic nucleus during functional mapping (purple). Spectrograms of the audio signals are shown. (**B**) Timing of behavioral events, relative to speech-onset. Heatmap of the duration of the auditory cue (AC) and speech production (SP) windows expressed as percentage across trials for each participant. Average phonetic accuracy of the produced syllables for each participant are shown on the right. (**C**) ECoG strips localizations. The coverage of the ECoG strips across participants is superimposed on three different target areas from the Destrieux atlas26: postcentral gyrus (purple), inferior frontal gyrus (blue) and superior temporal gyrus (green). Exemplary auditory-locked and speech-locked spectrograms of activity in the postcentral gyrus (purple sphere) and superior temporal gyrus (green sphere) after normalization with respect to the baseline are displayed. (**D**) MER localization. Coverage of single units across participants is depicted. Spheres denote the location of four exemplary neurons with different categories of instantaneous firing rate (IFR) modulation (expressed as Mean ± SEM percentage change for visualization): Increasing (red), Decreasing (blue), Mixed (green) and No (gray) firing rate modulation. (**E**) Exemplary transient spike-phase coupling in the a range during the speech production window (magenta dashed line). Spike timestamps, a oscillations, and instantaneous phase are illustrated. Magenta bars delineate the duration of the syllable triplet. List of abbreviations: ITI (intertrial interval) used as baseline, AC (auditory cue), SP (speech production) and IFR (instantaneous firing rate).

### Cortical potentials and subthalamic firing rates during speech production

Before addressing the complex interactions between cortical LFPs and STN single neuron firing during speech, we analyzed each of the signals independently using traditional methods. We decomposed cortical LFPs from lateral temporal and frontal cortex into time-frequency representations using Wavelet basis functions. We inspected LFP spectral components from 4 to 140 Hz. Expected cortical evoked activity was observed during both listening (locked to auditory cue onset) and speech production (locked to speech onset) (Figure 1) ^22–24^. Figure S2 illustrates different patterns of evoked activity in five representative electrodes from four participants. Spectrograms demonstrated consistent neural suppression in lower *α*–*β* frequencies (8-30 Hz) as well as elevation in the gamma *γ* range (50-150 Hz) during auditory cue presentation and speech production. A large fraction of STG electrodes displayed either transient or sustained increased *γ*–activity in response to auditory cues^25^. In line with previous work^13,24^, premotor and postcentral gyrus electrodes showed *β*–suppression and *γ*–elevation preceding the speech onset and during speech production. The same channels demonstrated above-baseline *β* increase (i.e., rebound) after the speech offset. Our data also revealed more complex response profiles, such as *γ*–activation of STG electrodes during speech (Figure S2), consistent with the role of STG during auditory feedback ^23,26^.

From STN microelectrode recordings, we identified 245 neurons. 211 were stable and isolated (Figure 1D shows recording density, on average 3.28 ± 1.25 neurons per recording session). Spike sorting and quality metrics were conducted as previously described ^14,20^. We found neurons’ instantaneous firing rates during the speech task was, as expected, heterogeneous both within and across recording session and patients (Figure 1D). N = 84/211 neurons (39.8%) exhibited a significant increase in their firing rate in a window around the speech onset. Other neurons (N = 37/211, 17.5%) displayed a decrease in their firing rate. Interestingly, N = 23/211 neurons (10.9%) showed mixed behavior with both increased and decreased firing rates. The remaining neurons (N = 67/211, 31.8%) did not exhibit a significant modulation of firing rate during the speech production task. For a comprehensive description of firing rate modulation in this dataset, see Lipski et al ^16^.

### STN neuronal spiking locks transiently in a specific frequency band with cortical field potentials

We then turned to the interactions between cortical LFPs and subcortical spikes. To investigate whether the subthalamic nucleus exhibits cortical phase-of-firing coding that is time-aligned with key events in speech production (Figure 1D), we employed a variable-window width spike-phase coupling (SPC) estimation procedure^18^. This method provides an unbiased, time-resolved estimate of the strength of spike-phase coupling across multiple frequencies (4-140 Hz) over the entire duration of the task, overcoming the limitations of traditional event-locked analyses that maximize the temporal precision only around the event of interest. To account for variability in the number of spikes during low and high firing rate periods, the method adjusts the window width centered around each computational bin, resulting in more accurate and less biased SPC estimates. The average window width was 0.15 ± 0.01 s with an average number of 350 ± 169 spikes across trials per window (see Methods for details). We obtained 19755 time-frequency SPC maps, with each map representing an STN neuron-cortical LFP pair. Maps specified SPC in frequency, from 4-140 Hz, and in time: from 0.75 s before the auditory cue onset to 0.75s after speech production termination. Neurons had on average 92.78 ± 33.28 cortical LFP pairs. Cluster-based permutation tests revealed that approximately 11% (2148/19755) of the time-frequency maps displayed significant SPC. The distribution of the percentage of significant SPC pairs across participants is illustrated in Figure S3.

When averaging only SPC maps (N = 2148) that were significant at the single-pair level, we observed SPC primarily in two distinct frequency ranges. STN neuron spiking uncoupled with cortical beta (13-21 Hz) with respect to baseline starting at auditory cue presentation and persisted throughout speech production (Figure 2A). After speech offset, STN spikes increased in β SPC beyond baseline, consistent with movement-offset β LFP power rebound ^27^. During the speech production interval, tonic (8-α)-SPC below 10 Hz became a prominent feature (Figure 2A). Notably, these results were consistent whether we averaged all SPC maps (N = 19755) or the most significant SPC map for each unit (N = 211) (Figure S4).

**Figure 2:**
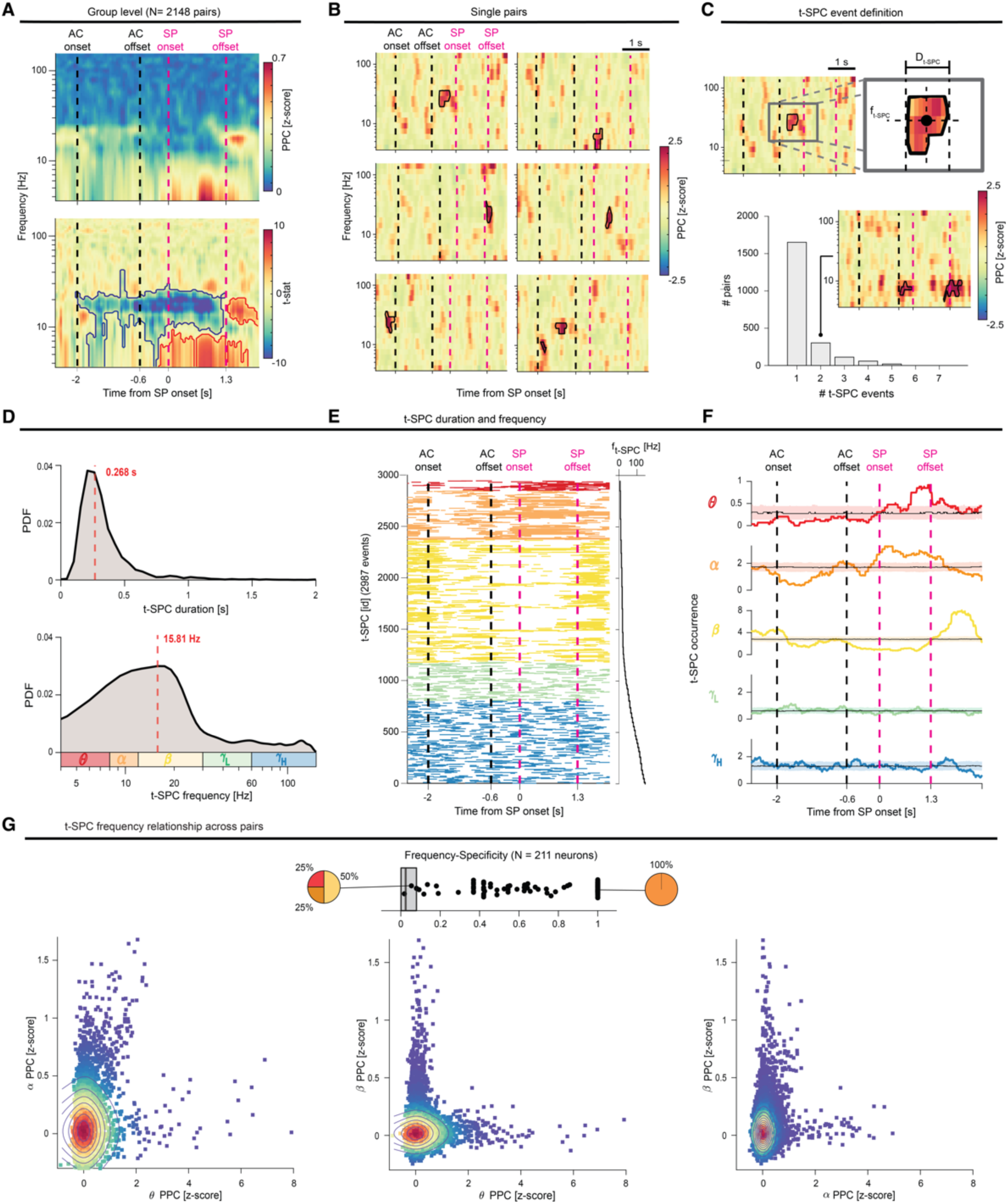
STN neurons show transient task-related coupling to the cortex in either θ-α or β bands. (**A**) Average of the spike-phase coupling (SPC) maps with significant spike-phase coupling (N = 2148 pairs). The pairwise-phase consistency (PPC) index is compared to the permutation distribution and expressed as z-score. Group-level statistical test (t-stat) of the significance of the z-score PPC with respect to the baseline across all the significant pairs. Red and blue lines contour regions of significant SPC increase or decrease, respectively. (**B**) Examples of single-pair SPC maps showing that STN neurons preferentially entrain cortical phases only during brief and transitory episodes. (**C**) Definition of the transient SPC event (t-SPC event) in a single-pair SPC map. We calculated the onset and offset times, temporal duration, and the frequency centroid for each t-SPC event. Most of pairs exhibit only one t-SPC event, as shown by the barplot. The inset plot depicts an exemplary SPC map with two t-SPC events in the same frequency band. (**D**) Distribution of the t-SPC duration and t-SPC frequency centroid. To augment the readability of the t-SPC frequency distribution, we adopted a logarithmic scale. Red dashed line depicts the median of the distribution. (**E**) List of the t-SPC events (N = 2987) ordered by frequency centroid. (**F**) t-SPC events occurrence grouped by frequency band. Shaded areas illustrate the 5th and 95th percentiles of the permutation distribution for the aggregation test. (**G**) (Top) The extent of the specificity of the t-SPC frequency band is depicted for each neuron (N = 211). The pie charts depict the proportion of t-SPC frequency band for two exemplary neurons. Dark gray boxes indicate the 5^th^ and 95^th^ percentile of the permutation distribution. (Bottom) Relationship between t-SPC spatial density (percentage of pairs with significant SPC, see Methods) in different frequency bands across pairs (left: θ vs α, center: θ vs β and right: α vs β). Colormap and contours indicate the 2D density of the scatter plot. Black and purple dashed lines denote auditory cue and speech production windows. Red dashed line depicts the median of the distribution. Black and purple dashed lines denote auditory cue and speech production windows. θ (red), α (dark orange), β (yellow), *γ*L (green) and *γ*H (blue).

We next sought to investigate the extent to which single-pair SPC maps accurately reflect the group-level SPC patterns observed during prolonged SPC changes. Strikingly, single-pair SPC maps showed that STN neurons entrain cortical LFP phases during transient episodes (Figure 2B), which we termed *t-SPC*. Although most significant SPC maps exhibit only one t-SPC (77%, 1652/2148), we also found examples with multiple (up to seven) t-SPC events, which can occur during different key events of the task and in different frequency bands (Figure 2B-C). Thus, periods of increased or suppressed group-level SPC reflect the type of t-SPC event most likely to occur.

We then characterized the task-related timing and frequency centroid of t-SPC events to test whether STN neurons display speech-related frequency-specific SPC. Our analysis revealed that t-SPC events had a median duration of 0.268 s and occurred most frequently in the beta range (∼16 Hz) (Figure 2D). We observed a mild negative correlation between t-SPC duration and t-SPC frequency centroid (R^2^ = 0.12 (ρ = −0.35), p < 0.001), suggesting that STN spiking is more likely to entrain for longer periods of time to lower frequencies. Figure 2E lists all the t-SPC events ordered by frequency band. 8-α t-SPC events significantly aggregate during speech production (p_perm_ < 0.05, permutation test) (Figure 2F). Notably, α t-SPC events occurred throughout the entire speech production duration, whereas θ t-SPC events signaled preferentially the final part of the utterance. Moreover, α t-SPC events decreased during the auditory cue presentation (p_perm_ < 0.05, permutation test). β t-SPC events were more prominent during the inter-trial interval (ITI), dipped during both auditory cue presentation and speech production, and rebounded above baseline levels after the termination of the utterance (p_perm_ < 0.05, permutation test). Neither *γ*_L_ nor *γ*_H_ t-SPC event occurrence showed prominent deviation from uniform distribution during the task.

SPC was highly specific within a particular frequency band at the single-pair level (Figure 2G). We employed an entropy-based metric to evaluate the frequency-specificity at the single neuron level. We then examined the relationship between neurons and frequency bands for neurons that had multiple t-SPC events: if a neuron coupled in one frequency band, was it more or less likely to couple in another band? A striking proportion of units (N = 203/211 96.2%, frequency-specificity ∼ 0.56) was significantly specific to a frequency band. Finally, neurons that exhibited increased (θ-α) t-SPC did not show any modulation of β SPC, and vice versa (Figure S5A-B).

We confirmed that t-SPC events observed in our data were not a by-product of the natural cluster tendency arising in small samples of random distribution. We evaluated the number of cycles of oscillations spanned by t-SPC events and the count of potential t-SPC events observed in the permuted SPC maps, previously used to convert the SPC maps into z-scores (see Methods). We used two cycles as the lower bound for a well-defined spike-phase coupling event, as is commonly chosen in local field potential oscillatory base analyses for beta bursts ^28,29^. 99.025% of t-SPC events had more than two cycles. There were approximately ten times more t-SPC events in the actual SPC maps compared to the shuffled SPC maps (p_perm_ < 0.001, permutation test) (Figure S6). These control analyses suggests that t-SPC events reflect genuine, physiological SPC mechanisms.

In summary, neurons tended to spike-phase couple to cortical LFPs transiently (for 0.25 s) and in a single frequency band. Neurons that did couple in multiple bands coupled in the θ and α range; these neurons seemed to couple to θ and α indiscriminately. Consequently, θ and α coupling were treated as a single entity in some analyses throughout the manuscript.

### Frequency dependence of preferred-phase of coupling reflects cortico-subthalamic time delays

The phase of a spike-phase coupling relationship has been shown to carry key information in other domains, such as working memory ^30,31^ and motor behavior ^18,20^. We tested whether STN neurons tended to spike at a preferred phase of the cortical LFP by quantifying the t-SPC phase and compared the consistency across t-SPC events over time. The polarity standardization procedure implemented in our pipeline allowed the comparison of phases across recordings (see Methods). We found that t-SPC events in the α range, but not other frequency bands, are locked around the same phase of firing (108°, during the decay after the peak of oscillation) during speech production (p< 0.05, Hodges-Ajne test) (Figure S7A). In contrast, β t-SPC events are uniformly locked around the trough (−90°) of the oscillation before and after the β t-SPC rebound (p< 0.05, Hodges-Ajne test), consistent with previous studies that employed electroencephalography or low-density EEG strips^21^.

We then utilized the t-SPC events in the 5-40Hz range to derive time information from the relationship between unwrapped t-SPC phase and t-SPC centroid frequency (see Methods). A positive slope indicates that STN spikes occur at a consistent time lag relative to the peak of the rhythmic ECoG activity, at a latency given by the slope divided by 2π. Grouping all data together, the gradient of the positive slope translated to a time lag of 40.92 ms between STN spikes and cortical activity (R^2^ = 0.76, p_perm_ < 0.001) (Figure S7B). When we performed the same analysis over time, we found that this relationship was particularly consistent during the β t-SPC rebound (39.81 ms, black dots in Figure S7B bottom). Interestingly, STN spiking led ECoG activity during speech production (−32 ms), while (8-α) t-SPC events were more frequent than β t-SPC events (green curve in Figure S7 bottom). This result might suggest that (8-α) t-SPC and β t-SPC observed here may reflect information flow between STN and cortex.

### STN-cortical spike-phase coupling during speech is temporally and topographically organized

Previous studies have shown that neural activity in the cortex and STN feature frequency-specific topographies during resting state^32,33^ and movement execution^34,35^. We next sought to delineate the frequency-specific topography of cortical t-SPC events during syllable repetition (Figure 3).

**Figure 3:**
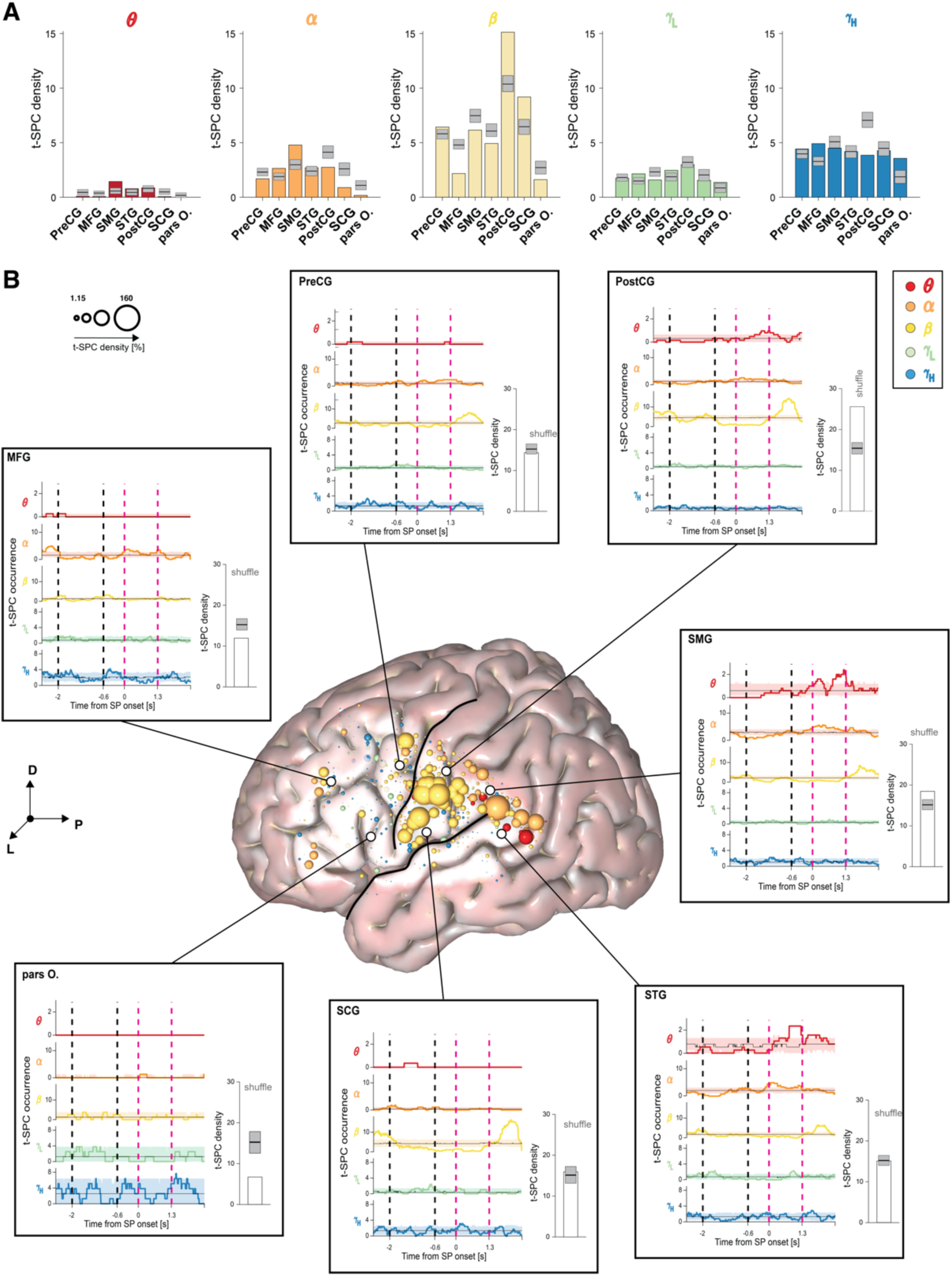
STN neurons couple to the SMG-pSTG in q-a during speech. (**A**) Spatial density of the t-SPC events across frequency bands in seven region of interests, as derived from the Destrieux atlas^26^. (**B**) Cortical spatial density map (radius of 2 mm) across frequency bands. The size of the spheres represents the degree to which t-SPC events are localized in a 2mm radius around the center of the spheres. Inset plots illustrate the overall t-SPC spatial density (white barplots) and t-SPC event occurrence in each region of interest. Shaded areas illustrate the 5^th^ and 95^th^ percentiles of the permutation distribution for the aggregation test. Dark gray boxes indicate the 5^th^ and 95^th^ percentile of the permutation distribution for the spatial preference test. Regions with spatial density higher or lower than the permutation distribution are labelled as high or low spatial preference. Black and purple dashed lines denote auditory cue and speech production windows. θ (red), α (dark orange), β (yellow), *γ*L (green) and *γ*H (blue). Black lines on the cortical surface delineate two anatomical landmarks: the Sylvain fissure (SF), which divides the temporal from the frontal and parietal lobes, and the Central sulcus (CS), which separates the Precentral gyrus (PreCG) anteriorly from the Postcentral gyrus (PostCG) posteriorly. List of cortical regions of interests: Precentral gyrus (PreCG), Postcentral gyrus (PostCG), Supramarginal gyrus (SMG), Subcentral gyrus (SCG), Superior temporal gyrus (STG), Middle frontal gyrus (MFG), and the orbital part of the inferior frontal gyrus (pars O.) The anatomical reference of frame shows the dorsal (D), lateral (L) and posterior (P) directions.

At the cortical level across all frequency bands, most t-SPC events were detected in the postcentral gyrus (PostCG, 17.13% significant pairs, t-SPC density = 25.57, p_perm_ < 0.05, permutation test) and supramarginal gyrus (SMG, 13.18% significant pairs, t-SPC density = 18.45, p_perm_ < 0.05, permutation test). Significant numbers of t-SPC events were found also in the superior temporal gyrus (STG, 13.74% significant pairs, t-SPC density = 15.06), precentral gyrus (PreCG, 10.74% significant pairs, t-SPC density = 14.09) and subcentral gyrus (SCG, 11.34% significant pairs, t-SPC density = 15.94), but t-SPC density was less frequent in the middle and inferior frontal areas (Figure 3A).

We observed aggregation (i.e., increased occurrence) of θ t-SPC events during speech production in SMG and STG (p_perm_ < 0.05). Similarly, α t-SPC events dispersed (i.e., reduced occurrence) during auditory cue presentation and aggregated during speech production in SMG and STG regions, in addition to other areas such as PreCG and PostCG (p_perm_ < 0.05). Baseline α t-SPC events were observed mostly in the MFG. Interestingly, α t-SPC events were not significantly spatially clustered around their centroid (x = −62.23 mm, y = −13.04 mm, z = 30.4 mm, p_perm_ = 1, Figure S8). β t-SPC events were present at baseline and later dispersed temporally from auditory cue presentation through speech production in the PostCG and SCG (p_perm_ < 0.05). Interestingly, the β t-SPC rebound was a more wide-spread phenomenon, observed in PostCG and SCG, and also in cortical regions like PreCG, SMG and STG which did not exhibit t-SPC during the baseline (p_perm_ < 0.05). β t-SPC events were spatially clustered around their centroid (x = −65.27 mm, y = −10.14 mm, z = 28.41 mm, p_perm_ < 0.05, Figure S8). *γ*_L_ and *γ*_H_ t-SPC events showed no preferential spatio-temporal distribution. We also compared the t-SPC event duration and centroid frequency across different region of interests. Longer t-SPC events with a lower frequency centroid occurred in the PostCG (∼320 ms, ∼20 Hz) and SMG (∼310 ms, ∼18 Hz) (p_perm_ < 0.05, permutation test) (Figure S9). These results indicate that different epochs of speech perception and production are accompanied by frequency-specific STN-cortical SPC signatures.

We next investigated the location of STN units involved in SPC. θ t-SPC events were significantly aggregated in the posterior-medial region of the STN (p_perm_ < 0.05, θ_1_ in Figure 4 and Figure S10). Moreover, θ t-SPC events were localized more dorsally (higher MNI z-coordinate) compared to t-SPC events in other frequency bands (Figure S10). Two α t-SPC hotspots (p_perm_ < 0.05, Figure 4 and Figure S10) were identified in the posterior-dorsal (α_1_) and posterior-ventral (α_2_) region of the STN. Of note, MFG SPC exclusively contributed to the posterior-ventral cluster. Overall, α t-SPC events were localized significantly inferior/ventrally (lower MNI-coordinate and PC2 coordinate) compared to t-SPC events in other frequency bands (Figure S11). Spatial density analysis in the β range demonstrated a more frequent emergence of β-SPC in the dorsolateral part of the STN during the baseline and rebound phases (p_perm_ < 0.05, β_1_ in Figure 4, Figure S7 and Figure S12C). β-SPC density appeared more focalized during rebound than during the baseline (Figure S11C). A transient increase in β-SPC events during auditory cue presentation was observed in the centro-medial region of the STN. Taken together, STN spikes during t-SPC events also revealed spatial patterns, albeit less distinct than for cortical LFPs. Table S2 summarizes centroids of t-SPC event location and peak of t-SPC spatial density for each frequency band on the cortex and STN.

**Figure 4:**
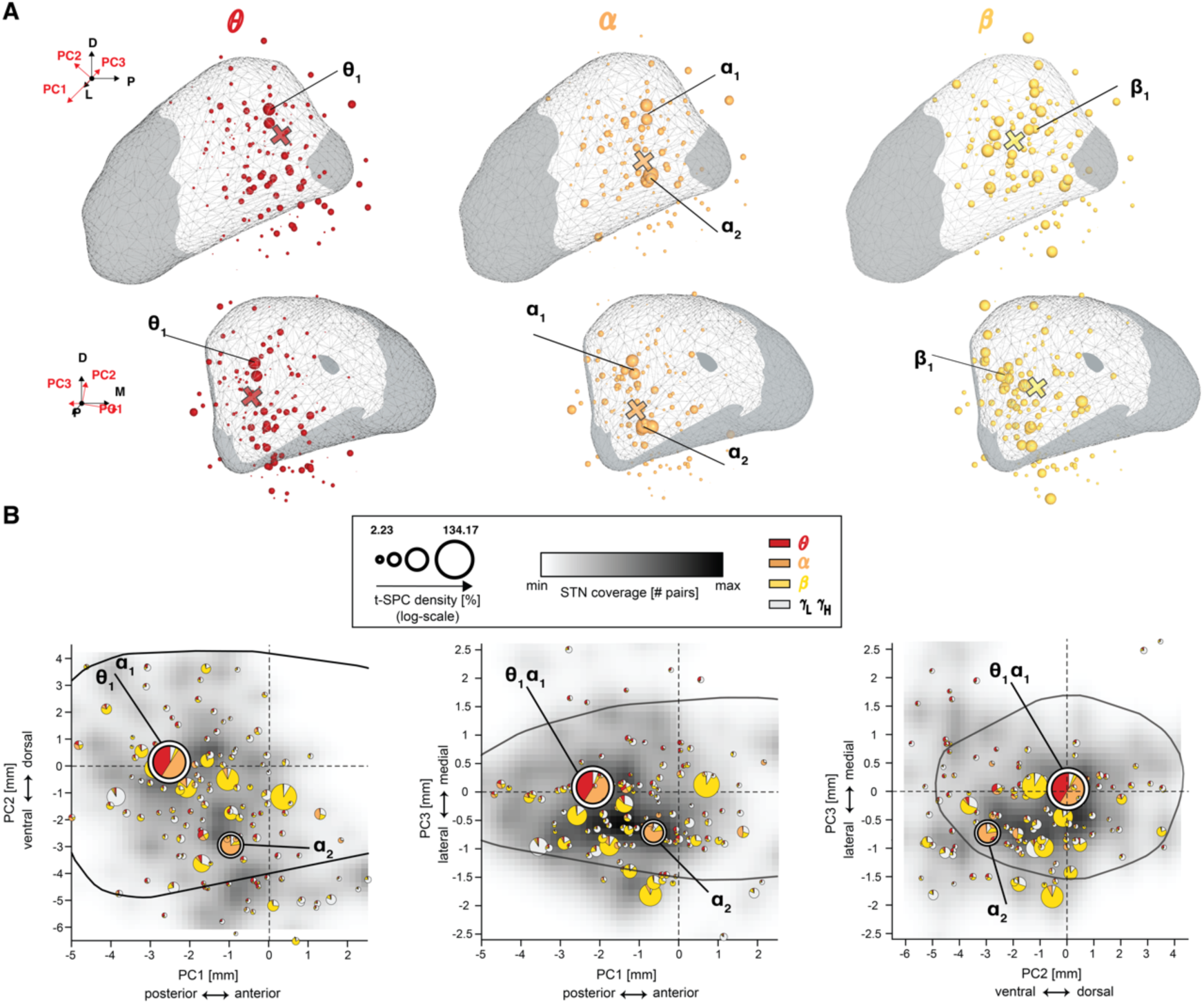
θ-α SPC neurons are localized in the posterior part of STN. (**A**) Subthalamic spatial density maps (radius of 1 mm) across frequency bands. STN regions sampled by microelectrode recordings are depicted as white overlay. Size of the spheres represents the degree to which t-SPC events are localized in a 2 mm radius around the center of each sphere. To augment the readability of the visualization, we adopted the logarithmic scale for the spatial density. The anatomical reference of frame shows the relative orientation between the dorsal (D), lateral (L) and posterior (P) directions and the first three principal components directions (PC1: anteriorposterior axis, PC2: dorso-ventral axis and PC3: medio-lateral). θ (red), α (dark orange) and β (light orange). Black and purple dashed lines denote auditory cue and speech production windows. Cross indicates the spatial centroid of the t-SPC event locations. θ_1_, α_1_, α_2_ and β_1_ depict the location of peaks of the t-SPC spatial density. (**B**) Spatial density of the t-SPC events mapped along the three principal component axes. The intersection of the two dashed black lines represents the STN center of mass. The radius of the pie-chart represents the t-SPC spatial density across bands. The black contour delineates the STN border as depicted by the DISTAL atlas. Note that principal component scores represent actual physical distances in mm. θ (red), α (dark orange), β (yellow).

### Firing rate modulation predicts preferred spike-phase coupling frequency

We used a variable-window width and pairwise-phase consistency (PPC) correction to ensure that changes in coupling strength were not merely a result of firing rate modulation. However, these two neural phenomena may represent distinct, yet overlapping, mechanisms of modulation^21,36^. To explore this possibility, we conducted a series of correlation analyses between t-SPC and firing rate across neurons (Figure S13). Importantly, we found no significant correlation between average firing rate and either average t-SPC density (R^2^ = 0.006, p_perm_ = 0.23) or t-SPC centroid frequency (R^2^ = 0.006, p_perm_ = 0.27).

To characterize the extent of overlap between changes in t-SPC density and firing rate modulation, we plotted SPC density changes (expressed as the difference) against firing rate modulation (expressed as z-score) with respect to the baseline (peak firing rate modulation) during speech production (see Methods) (Figure S13). We observed (8-α) t-SPC changes (increase: 31/211 units, 14.7% and decrease: 2/211 units, 0.9%) only in neurons exhibiting low or negative firing rate changes (< 5 z-score). Among the 27 neurons that displayed β t-SPC events during the baseline, 25 (92.6%) neurons significantly reduced their β t-SPC during speech production, either completely (22/25) or partially (3/25). Interestingly, a fraction of neurons (16/211, 7.6%) exhibited a slight increase in β t-SPC during speech production, suggesting a partial maintenance of the β SPC at the single-unit level. Importantly, we observed no significant differences in firing rate changes between neurons that decreased β t-SPC density during speech production or increased β t-SPC density after speech termination and neurons with no changes in β t-SPC density (Figure S13).

Comparing SPC across the different firing rate categories, we found that neurons whose firing rates were modulated by speech (18.51%) displayed a higher average t-SPC density compared to neurons without speech-related firing rate modulation (10.86%) (p_perm_ < 0.01, Figure S14). Among the speech-related neurons, those with decreased firing rate exhibited the highest average t-SPC density (23.43%), surpassing both increased (14.52%) and mixed firing rate modulation neurons (17.59%) (p_perm_ < 0.01). Both group-level SPC and t-SPC analyses indicated that only decreased firing rate neurons contributed significantly to the aggregation of θ t-SPC events during speech production (Figure S14). On the other hand, both decreased and increased firing rate neurons featured similar α t-SPC event occurrence profiles, occurring less frequently during auditory cue presentation and more frequently during speech production. Notably, increased, and mixed firing rate modulation neurons primarily contributed to the aggregation of β t-SPC events during the rebound phase, but only mixed firing rate modulation neurons displayed significant aggregation of β t-SPC events during the baseline. In contrast, no discernible pattern of t-SPC coupling emerged in neurons with no firing rate modulation across all levels of analysis (Figure S14).

We then examined changes in the centroid frequency and duration of t-SPC events across firing rate categories. Longer t-SPC events were observed in mixed firing rate neurons (∼0.31 s) and decreased firing rate neurons (∼0.31 s) with median centroid frequencies of 18 Hz and 20 Hz, respectively (Figure S15). When analyzed by firing rate category, only decreased firing rate neurons demonstrated a spatial preference for spike-phase coupling, which occurred with the SMG and STG, likely driven by t-SPC in the 8-α range. Together, these results suggest that while firing rate modulation alone does not fully explain the dynamics of SPC, the direction of firing rate change is correlated with distinctive patterns of SPC characterized by spectral, spatiotemporal, and anatomical features.

### Speech sound errors occur when θ-α spike-phase coupling is delayed

We considered spike-phase coupling as an indicator of information transfer between cortex and STN. We hypothesized that lower SPC might be correlated with speech performance defined as the phonetic accuracy, i.e., percentage of speech sound errors. We categorized trials into correct and error trials based on phonetic accuracy and computed SPC for each subset (see Figure 5). As high-frequency SPC did not show any significant task-related modulation, we restricted this analysis only in pairs with significant SPC in the 4 – 40 Hz range and at least 20 trials in each condition (827 pairs in 46 neurons). This analysis revealed that error trials exhibited lower θ-α SPC preceding speech production, followed by an overshoot in θ-α SPC after the speech termination (Figure 5A-B). Notably, no difference was observed during speech production. This trend held true whether we considered the SPC map (Figure 5A) or the t-SPC occurrence (Figure 5B) as a measure of SPC strength (p_perm_ < 0.05, permutation test). We also observed a significant increase of SPC in the β range in error trials but the result did not hold true when we looked at the SPC occurrence (compare Figure 5A with Figure 5B). We further hypothesized that θ-α t-SPC events occurred earlier in accurate trials than in error trials. We found that the median onset of θ-α t-SPC events is earlier in accurate trials than error trials, in a within-neuron analysis (p < 0.05, Figure 5C-D). t-SPC duration was not affected by phonetic accuracy before or after speech production (p > 0.05, Figure 5E).

**Figure 5:**
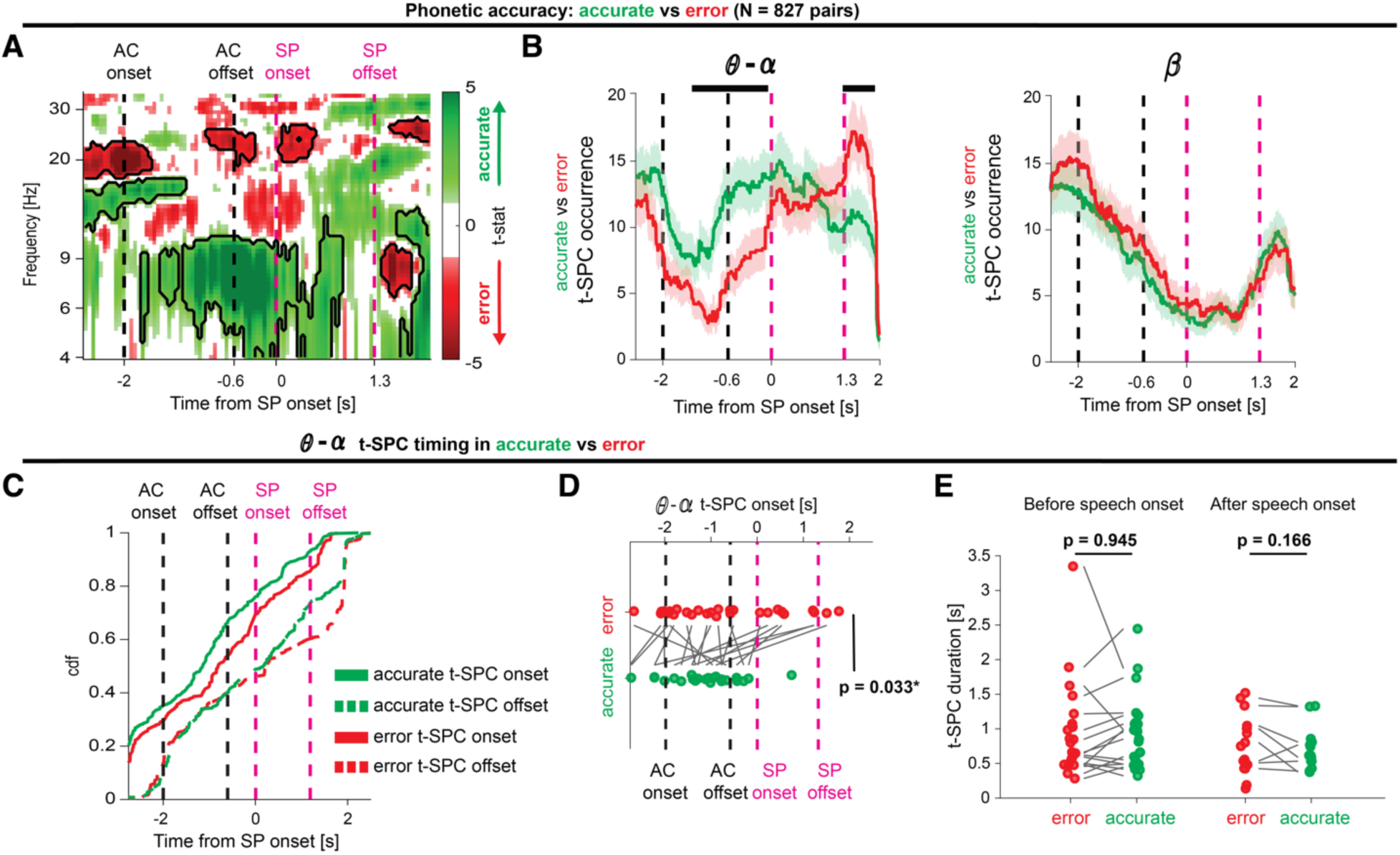
Phonetic errors are correlated with delayed θ-α STN-cortex coupling. (**A**) Comparison of the spike-phase coupling (SPC) maps (N = 827, 74 neurons in 18 participants) between trials with and without phonetic errors (see Methods). In error trials (red), θ spike-phase coupling before speech production onset was significantly lower than in correct trials (green). (**B**) t-SPC events occurrence in trials with and without phonetic errors grouped by frequency band (θ-α: left, β: right). Shaded areas illustrate the 5^th^ and 95^th^ percentiles of the bootstrapped distribution of t-SPC events occurrence. Black bars denote time bins in which error t-SPC occurrence is different between correct and error trials. (**C**) Cumulative distribution of the θ-α t-SPC onset and offset (see Figure 2C and Methods) in error and correct trials. (**D**) Comparison of the median θ-α t-SPC onset at the single neuron level (N = 20/46 neurons in 13 participants). (**E**) Comparison of the median θ-α t-SPC duration at the single neuron level before (N = 17/46 neurons in 13 participants) and after (N = 8/46 neurons in 13 participants) speech duration.

### Control analyses

To ensure the robustness and validity of our SPC estimates, we conducted a comprehensive set of control analyses (Figure S16-S17). First, we examined the impact of removing the event-related component of the ECoG signal before the computation of the SPC metric. After eliminating this component, the profile of the SPC maps remained largely unchanged, even at lower frequencies (Figure S16A-B). Indeed, we found no evidence of phase-reset of oscillations at the onset of auditory cue and speech production (Figure S16C). These findings suggest that the event-locked components (trial-averaged speech-locked signals) did not significantly influence the observed SPC pattern. Second, we assessed the influence of periods with high (top 10^th^ percentile) or low (low 10^th^ percentile) oscillation amplitude. When excluding these specific periods, the results remained comparable (Figure S17). These control analyses reinforce the significance of the observed SPC patterns.

## Discussion

In this study, we found robust evidence of transient (∼268 ms) spike-phase coupling events that occur between STN neurons and broad cortical regions during a syllable triplet repetition task. We found that cortical potentials and subthalamic neurons exhibit complex dynamics during speech, organized both, temporally and topographically. Events of spike-phase coupling in theta and alpha range mainly dominated within auditory-sensorimotor integration areas through the posterior perisylvian cortex. This region contains the “dorsal stream” ^37^, i.e., STG and SMG, and SPC onsets here occurred earlier before phonetically accurate speech production trials. This finding suggests that STN neurons participate in the implementation of speech motor programs to minimize the occurrence of speech sound errors.

Our results align with the notion of the cortico-basal ganglia thalamic loop subserving temporal-integration for modulating motor control ^38–40^, consistent with previous evidence of SPC between STN neurons and cortical field potentials during limb movement ^19–21,36,41^. Similarly, cortical oscillations manifest as transient bursts, whose onset is preceded by an increase of spike-phase coupling with the STN^42^. These transient periods might represent “open windows” for effective communication between STN and cortex. The duration of these windows may be constrained by a slower subcortical neural timescale and requirements of a given motor instantiation^42,43^.

Our observations suggest that the contribution of θ-α SPC between the STN and the posterior opercular cortex may arise at multiple stages of speech production, including phonological processing, executive processing and articulatory processing ^44^. Prior studies have reported that STN neurons encode phonetic characteristics during speech production ^16^ and that STG lexical-encoding gamma signals are projected into the STN prior to speech production ^17^. A recent study showed that changes in functional connectivity between STN and language regions predicted the downstream effect of dopaminergic medication on speech-related cognitive performance ^45^. Coming from a different recording modality and different measure of connectivity, our results bolster the finding that STN is involved in speech circuitry. θ-α SPC occurred earlier before speech onset when participants accurately pronounce the target phoneme, in contrast to trials with phonetic errors, their timing was later, and magnitude was less before speech but overshot at termination. Additionally, alpha SPC was present in other cortical regions, including the PostCG and PreCG, as well as the STG and SMG, suggesting its role in internal monitoring of speech output and providing feedback on its match (or mismatch) with the expected auditory-sensorimotor prediction. This hypothesis for STN SPC function complements previous studies describing gamma amplitude changes in the STG and SMG ^23,46^ and STN single-unit correlates in speech ^13,14,16,17^, reinforcing the importance of the STN as a hub that processes multimodal cortical information ^47^.

At the neuron level, changes in spike-phase coupling during speech production were not correlated with speech-related changes in the instantaneous firing rate or the baseline instantaneous firing rate. Other studies have reported similar decoupling between SPC and firing rate in STN neurons during movement ^19,20^. Moreover, we found that neurons fired at comparable intensity, irrespective of their significant SPC display, except for theta SPC events. These were predominantly driven by neurons that decreased their firing rate in the early stages of speech production. Interestingly, we previously found that neurons often reducing their firing rates during speech did so in association with the onset of the auditory cue ^14^. Our findings underscore the notion that information can traverse the cortico-basal ganglia loop either through changes in the firing rate activity or spike timing. We also found that the pronounced overall reduction of beta SPC observed at the population level during speech production did not reflect a uniform reduction of SPC at the single-unit level. A small subset of neurons (7.6%) increased their beta SPC during speech production, suggesting a partial maintained beta SPC and the presence of a distinct functional beta SPC network. This aligns with two other studies that reported similar subpopulations of STN neurons, which increased their beta SPC during motor activity ^20,21^. The functional relevance of this partially maintained beta SPC during speech production remains uncertain.

Consistent with other studies, we showed that cortical oscillations lead in phase over firing activity in the STN ^19–21^, especially in the beta range post-speech. We found a delay of ∼41 ms aligning closely with other reports ^20,21^. Notably, the STN led the cortex exclusively when SPC in theta and alpha was more pronounced during speech production. While this delay does not necessarily signify synaptic transmission delay, it is consistent with the transmission of information through the cortico-basal ganglia loop. In light of our recent findings suggesting the presence of monosynaptic connections between non-motor regions of the cortex (sensorimotor and auditory areas) and the STN ^9^, it is not out of the question that the SPC we report here is anatomically rooted in the hyperdirect pathway.

Our results have important implications for theoretical frameworks of both basal ganglia function and speech production. These findings illustrate how classic firing-rate based models are insufficient for explaining basal ganglia circuit behavior. In the context of speech production, models such as DIVA^5^ and SFC^6^, are predominantly cortico-centric and do not explicitly incorporate relevant basal ganglia nodes such as STN.

Beyond expanding theoretical frameworks, our results may have important implications for clinical therapies. Although many Parkinsonian motor symptoms can often be satisfactorily controlled by STN DBS, stimulation-induced effects on the speech motor system can be heterogeneous ^48^. How can stimulating the same target nucleus consistently ameliorate some parkinsonian symptoms yet have mixed and variable effects on the speech motor system? Our results align with the notion that variability in DBS lead placement can explain most of the reported variance of outcomes in the literature ^32^. Relative to the optimal therapeutic target defined by Caire et al ^49^(x = −12.6 mm, y = −13.4 mm, z = −5.9 mm), the spatial centroid of our speech-related STN SPC topographies is at least 2.5 mm distant and overall located more posterior and ventral (Table S2). This aligns with studies that found detrimental effects on speech outcomes when stimulating more posteriorly ^50–53^ and ventrally ^54^. However, all these studies simply compare the speech outcomes between DBS ON and DBS OFF conditions without considering the stimulation amplitude and the spread of the stimulation volume towards neighboring regions. Hypotheses for future investigation include stimulating the STN in areas of SPC density peaks to test for altered integration of sensorimotor and auditory signals.

Our findings should be interpreted in the light of several limitations. First, a recent study suggested that spike-phase coupling does not mean that STN neurons generate and resonate rhythms coherent with the cortical synaptic input activity or that cortico-subcortical coherence has a functional effect on information transmission gain ^55^. Consequently, spike-phase coupling may simply reflect the fact that STN neurons integrate cortical inputs to some extent. Analysis of spike-phase coupling in both directions and causal investigations are necessary to circumvent these theoretical limitations ^55,56^. Second, caution is necessary when interpreting SPC results, as the presence of SPC between STN neurons and narrowband cortical oscillations does not imply that STN neurons generate and resonate coherent rhythms with the cortex ^55^. For example, neurons displaying significant theta SPC with the STG do not necessarily oscillate in the theta-rhythm at the population level. Therefore, our frequency-wise STN SPC topographies during speech production would not necessarily align with STN power-based topographies based on local field potentials recorded at rest ^33,57^. Third, phase-based time delays do not necessarily reflect synaptic transmission delays ^58^, but likely reflect the combination of cortical and subcortical inputs to the STN that become transiently stable ^21,41^. Fourth, ECoG coverage varied across participants and spanned a limited region of the cortical surface. However, we were able to perform group-level analysis by considering only regions of interest with adequate coverage. We cannot rule out any other interaction of the STN with other cortical regions. Fifth, we were unable to record single-unit activity across the entire volume of the STN due to unequal spatial sampling. Most microelectrode trajectories traversed the dorsolateral part of the STN, the clinical target for PD DBS ^32^. Hence, sampling of the ventro-medial region of the STN is limited. Differences in cortical coverage and subcortical sampling confounded correlative analyses between neural signals and clinical scales. Sixth, our findings are based on Parkinson’s Disease patients and caution must be exercised when interpretations of human neurophysiology are drawn from observations collected in a pathological state. Specifically, differences in the STN baseline firing rate ^59^ and abnormal subcortical beta oscillations ^29,60–62^ that characterize the Parkinsonian state may confound distinction of whether our observations generalize to speech in individuals without PD.

In summary, we discovered evidence that STN neurons are linked to the phase of the cortical oscillations during speech. These insights provide a deeper understanding of how different types of information are processed in basal ganglia-cortical loops and have significant implications for understanding the role of the human basal ganglia in sensorimotor integration for speech and other behaviors^63^.

## Methods

### Participants

Electrophysiological signals were recorded intraoperatively from 24 participants (20 males and 4 females, age: 65.4 ± 7.1 years) with Parkinson’s Disease undergoing awake stereotactic neurosurgery for implantation of DBS electrodes in the subthalamic nucleus (Table S1 for clinical details). Participants performed up to 4 sessions of the task, leading to a total of 64 sessions, after overnight dopaminergic medication withdrawal. All procedures were approved by the University of Pittsburgh Institutional Review Board (IRB Protocol #PRO13110420) and all participants provided informed consent to participate in the study.

### Method details

#### Speech production task

Participants were tasked to intraoperatively repeat aloud consonant-vowel (CV) syllable triplets. The stimuli were presented auditorily via earphones (Etymotic ER-4 with ER38-14F Foam Eartips) and were delivered at either low (∼50 dB SPL) or high (∼70 dB SPL) volume using BCI2000 as stimulus presentation software. The absolute intensity was tailored to each participant’s comfort level, keeping fixed the difference between high and low conditions at 25 dB SPL. The experiment utilized a set of phonemes consisting of four consonants (/v/, /t/, /s/, /g/) with different manners of articulation and three cardinal vowels (/i/, /a/, /u/) with distinctive acoustic properties. We created a unique set of 120 triplets of CV syllables, forbidding CV repetition within the triplet and balancing syllables and phoneme occurrence, and CV position within the triplet across a run of the task. The audio produced by the participant was recorded with an PRM1 Microphone (PreSonus Audio Electronics Inc., Baton Rouge, LA, USA) at 96kHz using the Zoom-H6 portable audio recorder (Zoom Corp., Hauppauge, NY, USA).

#### Neural recordings

As part of the standard DBS clinical procedure, functional mapping of the STN was performed using microelectrode recordings (MER) acquired with the Neuro-Omega recording system (Alpha-Omega Engineering, Nof HaGalil, Israel) using parylene insulated tungsten microelectrodes (25 μm in diameter, 100 μm in length). The microelectrodes were oriented using three trajectories (Central, Posterior and Medial) of a standard cross-shaped Ben-Gun array with a 2 mm center-to-center shaping. MER signals were referenced to the metal screw holding one of the guide cannulas used to carry the microelectrodes and recorded at 44 KHz. Prior to STN mapping, participants were temporarily implanted with two high-density subdural electrocorticography (ECoG) strips consisting of 54 or 63 contacts, respectively (PMT Cortact). These strips were placed through the standard burr hole, targeting the left ventral sensorimotor cortex, and left inferior frontal gyrus. Signals from ECoG contacts were referenced to a sterile stainless-steel subdermal needle electrode placed on the scalp and acquired at 30kHz with a Grapevine Neural Interface Processor equipped with Micro2 Front Ends (Ripple LLC, Salt Lake City, UT, USA).

#### Electrode localization

We localized the ECoG strips and DBS leads using well-established pipelines in the literature. For ECoG strips, contact locations were determined using the Randazzo localization method^64^ that utilizes a preoperative T1 weighted MRI scan, an intraoperative fluoroscopy and a postoperative CT scan (github.com/Brain-Modulation-Lab/ECoG_localization). CT and MRI were coregistered using SPM (https://www.fil.ion.ucl.ac.uk/spm/) and then rendered into a 3D skull and brain using Osirix (www.osirix-viewer.com) and Freesurfer (https://surfer.nmr.mgh.harvard.edu) software. The position of the frame’s tips on the skull and the implanted DBS leads were used as fiducial markers, which were coregistered and aligned with the projection observed in the fluoroscopy. The position of the contacts in the ECoG strip were manually marked on the fluoroscopy image and then projected to the convex hull of the cortical surface. To extract native coordinates of individual contacts, we leveraged the known layout of the ECoG strip. All coordinates were then transformed into the ICBM MNI152 Non-Linear Asymmetric 2009b space, employing the Symmetric Diffeomorphism algorithm implemented in Advanced Normalization Tools (ATNs). For DBS lead reconstruction, we used the Lead-DBS localization pipeline^65^. Briefly, the process involved coregistering the MRI and CT scans, and manually identifying the position of individual contacts based on the CT artifact, constrained by the geometry of the DBS lead used. The coordinates for the leads in each participant’s native space were rendered after this process. Custom Matlab scripts (github.com/Brain-Modulation-Lab/Lead_MER) were then used to calculate the position of the micro- and macro-recordings from the functional mapping based on the position of the lead, the known depth, and tract along which the lead was implanted in each hemisphere. Anatomical labels were assigned to each contact based on the Destrieux atlas^66^ for cortical contacts, and the DISTAL atlas^67^ for subcortical contacts.

### Quantification and statistical analysis

#### Phonetic coding

To extract phoneme characteristics from the produced speech signals such as onset and offset times, IPA code and accuracy, we employed a custom Matlab GUI (github.com/Brain-Modulation-Lab/SpeechCodingApp). Phonetic coding of each produced phoneme was performed by a trained team of speech pathology students using Praat (https://www.fon.hum.uva.nl/praat/).Discrepancies between produced phoneme and target phoneme were labelled as phonetic errors. We identified three types of errors: consonant substitution (e.g., */g/* produced as */v/*), vowel substitution (e.g., */u/* produced as */i/*) and phonemic omission (e.g., */su/ /ti/ /ga/* produced as */su/ /i/ /ga/*).

#### Behavioral events

For each trial, we defined four different behavioral epochs: baseline epoch as a 500 ms time window between −550 ms and −50 ms prior to the auditory cue onset, auditory cue presentation as the window during which syllable triplets were presented auditorily (∼1.5 s duration), speech production as the variable time window during which participants repeated aloud the syllable triplet (∼1.6 s duration on average) and post-speech as the 500 ms time window after the speech offset.

#### Electrophysiological data preprocessing

ECoG preprocessing was performed using custom code based on the Fieldtrip toolbox^68^ implemented in Matlab, available at (github.com/Brain-Modulation-Lab/bml). To temporarily align recordings from the Ripple, Neuro-Omega and Zoom-H6 systems, we employed a linear time-warping algorithm based on the stimulus and produced audio channels. The alignment was continuous throughout the entire recording session with sub-millimetric precision. Data was low pass filtered at 250Hz using a 4^th^ order Butterworth filter, downsampled to 1KHz and stored as a Fieldtrip object. Metadata such as descriptions of each session, phonetic coding, event times and electrode locations were stored in annotation tables. We applied a 5^th^ order high-pass Butterworth filter at 1 Hz to remove drifts and low-frequency components. Segments with conspicuous high-power artifacts were identified using an automatic data cleaning procedure^69^, based on a power-based threshold. Specifically, we extracted power at frequencies in different canonical bands (3 Hz for δ, 6 Hz for θ, 10 Hz for α, 21 Hz for β, 45 Hz for γ_L_ and 160 Hz for γ_H_) by convolving ECoG signals with a 9-cycles Morlet wavelet. A time bin was classified as artifactual if its log-transformed power in any band exceeded a threshold defined as the mean ± 2.5 std (∼10-fold higher the mean). Trials with time segments flagged as artifactual were discarded and channels with more than 30% of artifactual time bins were not included in the analysis.

#### Spike sorting

Spike sorting was performed using Plexon (https://plexon.com/products/offline-sorter/) as previously described^14^. We used a 4^th^ order Butterworth high-pass filter with a cutoff frequency at 200 Hz and set a manual threshold to extract putative waveforms. Single units were discriminated and graded based on factors such as cluster isolation in the principal component, the spike sorting’s stability over time, a refractory period of at least 3 ms in the inter-spike interval distribution, and the shape of the waveform.

#### Instantaneous firing rate

To analyze changes in spike rate activity, we followed the procedure described in ^14^. We sought elevated and reduced firing activity by computing the instantaneous firing rate (gaussian kernel, σ = 25 ms) and the inter-spike interval (smoothing window 25 ms), which scales with the reciprocal of the instantaneous firing rate, respectively. We aligned these quantities with speech production onset and analyzed time bins from auditory cue onset through speech production offset. A neuron was considered as Decreasing firing rate neuron if the inter-spike interval exceeded for at least 100 ms the upper 5% of a normal distribution with mean and standard deviation calculated during the baseline period. Similarly, a neuron was considered as Increasing firing rate neuron if the instantaneous firing rate exceeded for at least 100 ms the upper 5% of a normal distribution with mean and standard deviation calculated during the baseline period. Neurons that exhibit both modulations were named as Mixed firing rate modulation neurons, while neurons that did not exhibit any significant speech-related firing changes were labelled as No firing rate modulation neurons. For a comprehensive description of the firing rate modulation, please refer to Lipski et al ^16^.

#### Time-frequency decomposition

Time-varying power and phase were obtained by applying the Hilbert Transform to the band-pass filtered ECoG signal. The signal was bandpass filtered using a 4^th^ order Butterworth filter, with the following frequency ranges: 5–8 Hz for theta, 8–12 Hz for alpha, 12–20 Hz for low beta, 20– 30 Hz for high beta, and center frequencies ranging from 40 to 150 Hz with a bin width of 10 Hz, incrementing by 10 Hz.

#### Spike-phase coupling implementation

To calculate spike-phase coupling, we considered each possible pair of neurons and ECoG signals that were synchronously recorded. We enforced the following criterion for determining the eligibility of pairs (N = 19755) for subsequent analysis: a minimum 10 trials with stable firing rate and clean ECoG signal. The strength of the spike-phase coupling was quantified by the phase-locking value (PLV), which represents the magnitude of the circular average of unit complex vectors corresponding to the ECoG phase at the time of each spike φ_t_, as follows:

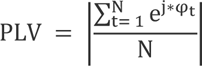

where N is the number of spikes included in the window. PLV is bounded between 0 and 1, indicating lack or perfect spike-phase coupling, respectively. Importantly, PLV is inflated toward 1 when N is low. When N is sufficiently large (N > 50), the pairwise phase consistency (PPC) yields an unbiased estimator of spike-phase coupling^70^, as follows:

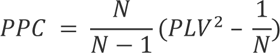

In the absence of spike-phase coupling, PPC is expected to be centered around zero, including negative value, when N is finite. As N increases towards infinity, the PPC tends to *PLV*^2^. To ensure that changes in spike-phase coupling depicted genuine and comparable neural correlates, methodological considerations must be discussed. First, the presence of speech-related fluctuations in the instantaneous firing rate poses a challenge in selecting a fixed window size for calculating the phase-locking value (PLV) or PPC over time. This is because variable N can result in uncontrollable and variables biases. Furthermore, low N can lead to noisy estimates of PPC. Second, intra- and inter-participant variability in speech production onset and duration makes event-locked analysis less accurate for the alignment of data around all key task events and not just for one single event (the one used for locking). To overcome all these limitations, we employed a variable-window width SPC estimation procedure developed by Fischer and colleagues^18^. First, we defined five intervals: from 0.75 before auditory cue to auditory cue onset, from auditory cue onset to offset, from auditory cue offset to speech production onset, from speech production onset to offset, and from speech production offset to 0.75 s after. Second, we subdivided these intervals into 8 equidistant anchor points, resulting in 36 anchor points for each trial. Third, we scaled the width of the window centered at each anchor point such that the sum of spikes (N) across all trials would match a target number as closely as possible. The target number was defined as the average number of spikes in a window of 0.15 s, but always greater than 25 to avoid fewer representative samples. This process allowed to enlarge/shrink the computational window during reduced/increased firing rate periods, ensuring that N remained constant over time. Note that we allowed variable number of spikes across participants to reduce variability in the window width. Finally, each window was placed symmetrically around each anchor point, and we subsequently calculated the PLV metric and applied the PPC correction. The resulting 19755 SPC maps (PPC values of size 16 frequency bins x 36 time points) were smoothed using a time-frequency window ([2, 2] size) and rescaled to the average duration of the event intervals. These maps were then event-locked and averaged across pairs and participants.

#### Phase polarity standardization

When computing the PLV or PPC, information about the preferred phase is not retained. To identify the preferred phase at which spikes are bundled, we calculated the circular mean using the CircStat toolbox^71^. However, it is important to exercise caution when comparing preferred phases across recordings due to the relative orientation between neural sources and electrodes (i.e. source mixing) and the use of different re-referencing schemas, as these factors can obscure the interpretation of the instantaneous absolute phase^72^. For instance, by applying the bipolar schema, the order of subtraction between two electrodes can flip throughs to peaks and peaks to throughs. To ensure that phases were meaningfully computed across recordings, we applied an automatized polarity-standardization procedure^18^. Specifically, we flipped phases (+ π) such that gamma peaks in the 60-80 Hz range consistently coincided with increases in the local high-frequency activity, which served as a polarity-invariant proxy of background unit activity^73,74^. We computed this proxy by high-pass filtering the ECoG signal at 300 Hz, full-wave rectifying it, and low-pass filtering it with a cut-off of 100 Hz. Flipping procedure was required in 7111/19755 pairs (36%).

#### Spike-phase coupling events

To further correct the SPC maps for any residual bias and identify genuine increases in SPC, we converted PPC values into z-scores relative to a permutation distribution and performed a cluster-based permutation test^75^. We built the permutation distribution by shuffling the trial-association between STN spikes and EcoG phases 500 times. We paired spike timings from the i-th trial with EcoG phases from the j-th trial (where I ≠ j). Importantly, to be conservative and preserve the natural appearance of clusters, we applied the same randomization across time-frequency bins. Suprathreshold clusters (p < 0.05) were identified in both the original SPC map and in each permutation SPC map by computing the z-score relative to the permutation distribution. If the absolute sum of the z-scores within the original suprathreshold clusters exceeded the 95^th^ percentile of the 500 largest absolute sums of z-scores from the permutation distribution, it was considered statistically significant. These significant clusters in the SPC map were referred to as transient-SPC events (t-SPC events). SPC maps that contained at least one t-SPC event were considered significant.

#### Spike-phase coupling event characteristics

To fully characterize each t-SPC event, we defined a set of characteristics in the time, frequency, and phase domain. In the temporal domain, we calculated the onset and offset times, temporal duration, and the center of the event (i.e. the mean of the onset and offset). For the frequency domain, we calculated the frequency centroid, as follows:

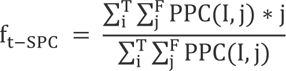

where I and j are the i-th time bin and j-th frequency bin enclosed within the boundaries of the t-SPC event. In the phase domain, we calculated the circular mean of the t-SPC event phases.

#### Time occurrence

We aimed to quantify the temporal occurrence of t-SPC events, which we defined as the likelihood of observing at least one t-SPC event in a given STN neuron-ECoG contact pair during each time bin and in each frequency band. To achieve this, we transformed each t-SPC event into a binarized vector, where each time-point (at intervals of 5 ms) was labeled either as part of a t-SPC event and assigned a value of 1, or as part of a non-t-SPC period and assigned a value of 0. We then calculated the mean of these binarized vectors at each time bin, expressed as a percentage. Higher values of this quantity indicated a greater temporal aggregation (i.e., overlap) of t-SPC over time, whereas lower values indicated dispersion. To identify significant time windows of aggregation or dispersion, we employed a permutation test, adapting the approach used to calculate significant beta bursts overlap, as described in ^28^. We generated a permutation distribution of the time occurrence due to chance by setting a variable break point in the 0s of the binarized vectors (no slicing of t-SPC events), reversing the two segments, and joining them together. We repeated this process 500 times and extracted the permutation distribution over time. We considered t-SPC events to be significantly dispersed when the time occurrence fell below the 5^th^ percentile of the permutation distribution, and significantly aggregated when the time occurrence rose above the 95^th^ percentile of the permutation distribution. By applying this method, we were able to rigorously determine changes in time occurrence even when the number of t-SPC events was low, and to eliminate spurious trends of aggregation when the number of t-SPC events was high.

#### Spatial density

We quantified the spatial density of t-SPC events by calculating the ratio between the number of t-SPC events and pairs and expressing it as a percentage, both at the cortical and STN level. To calculate the spatial density on the cortical surface, we used two region-of-interest-based methods. In the first method, we identified 7 regions of interest in the Destrieux atlas^66^ that satisfied a minimum coverage criterion (>7 participants and >100 pairs): Precentral gyrus (PreCG), Postcentral gyrus (PostCG), Supramarginal gyrus (SMG), Superior temporal gyrus (STG), Middle frontal gyrus (MFG), and the orbital part of the inferior frontal gyrus (pars O.). We calculated the spatial density in each region of interest and determined whether t-SPC events were preferentially located in any region of the brain or whether they exhibited no spatial preference, both overall and within frequency band. To test for spatial preference, we created a null permutation distribution (spatial uniform distribution) by shuffling the spatial label of each t-SPC event 500 times. We then compared the spatial density of the original data to the 5^th^-95^th^ percentiles of the spatial density permutation distribution. Regions with spatial density below the 5^th^ percentile and above the 95^th^ percentile were classified as having “low” or “high” spatial preference, respectively. In the second method, we created a cortical spatial density map by calculating the spatial density in a spherical region of interest with a radius of 2 mm centered around the ECoG recording locations in the MNI space. These maps were converted and displayed in SurfIce as nodes. For the STN domain, we built an STN spatial density map by locating spheres (1 mm radius) around the STN neuron locations. Again, we used SurfIce for visualization. For each spatial density map, we identified the peak in each frequency band. Additionally, we projected the STN neurons locations onto the three principal directions of the STN extracted from the DISTAL atlas image^67^, following the procedure as described in ^57^. To preserve the physical meaning, i.e., distance in mm, of the principal component decomposition, we multiplied the principal component scores by the standard deviation of the MNI coordinates. The principal component coordinates (PC1: antero-posterior direction, PC2: dorso-ventral direction and PC3: medio-lateral direction) represents a more suitable reference of frame, as the STN is not fully spatially aligned with the MNI coordinates (Figure S11A-B). Spatial density computation was repeated for each of the three principal axes using a side of 0.8 mm. We also employed the same formula to calculate the spatial density at the STN neuron level (ratio between the number of t-SPC events and pairs in each neuron). We used this level of analysis to compare SPC at different epochs of the task or frequency bands across neurons and to control for the effect of the firing rate (see **Control analysis**). When comparing β t-SPC during baseline and speech production epoch, we included in the computation only those t-SPC events with mean time in the related behavioral epoch. We also assessed the extent to which neuron preferentially couple to the same frequency band. We normalized the spatial density across frequency bands (total sum = 1) and defined the frequency-specificity as one minus the entropy of the normalized distribution. With this definition, high (e.g., peaked distribution) and low specificity (e.g., uniform distribution) are mapped onto 1 and 0 values, respectively.

#### Spatial aggregation

To extend and further corroborate our findings in the spatial domain, we also conducted a region-of-interest-free analysis, both at the cortical and STN level. MNI and PC coordinates (and their centroid) of t-SPC events were compared across frequency bands using a permutation test. We then investigated whether t-SPC event locations (within each frequency band) were more spatially aggregated around their centroid than expected by chance (uniform distribution). To this end, we computed the average Euclidean distance between t-SPC events locations and their centroid^57^, and compared against a null distribution of surrogate average Euclidean distances obtained by randomly sampling recording locations 500 times.

#### Relationship between STN spike-phase-coupling topographies and DBS anatomical STN targets

To investigate the relationship between the frequency-specific STN topographies and optimal DBS target for motor symptoms control in PD, we calculated the Euclidean distance between frequency-wise spatial centroids and location of DBS contacts commonly used for therapeutic stimulation^32,49^.

#### Time delay analysis

As STN neurons often lock to cortical signals within a narrow frequency range, power-based estimates of time delay between STN and cortex might be suboptimal^1^. We calculated time delays using the phase-based analysis, as described in ^21^. First, we computed the mean preferred phase of units that were significantly locked in each frequency bin (5-30 Hz range). We then averaged these phases to obtain a grand average phase for each frequency band. By analyzing the gradient of these phases, we determined whether the ECoG channel led (positive sign) or lagged (negative sign) relative to STN neuron, and at what latency this occurred. To test the significance of the time delay, we repeated 500 times the computation using randomly selecting mean angles from each frequency bin. To obtain a p-value, we compared the correlation coefficient in the original data and the 5^th^-95^th^ percentiles of the correlation coefficient permutation distribution.

#### Relationship between spike-phase coupling and speech behavior

To examine the link between spike-phase coupling and speech behavior, two sets of analyses were conducted on phonetic accuracy. We restricted this analysis only in the low-frequency range (4-40 Hz). For phonetic accuracy, trials were categorized into correct (100%) and error (< 100%) groups. Only significant (from the main analysis) pairs with ≥20 trials in each condition were included (46 neurons, 827 pairs across 18 participants).

To balance the number of trials, we subsampled 20 trials in each condition and run SPC pipeline 50 times. SPC maps were subtracted and averaged across subsamples. We converted the PPC values into z-scores relative to a permutation distribution defined as the difference between the permuted values in the two conditions. Significance was evaluated using the same procedure as above (see **Spike-phase coupling events**). The significant clusters in the difference SPC map were referred as t-SPC events signaling time-frequency bins in which the first condition was either higher or lower than the second one according to the sign of the z-value. For statistical comparison at the group level between the two conditions, we converted the z-scores to t-values and generated 500 permuted samples by randomly permuting the order of subtraction of the two SPC maps. P-values were estimated using the null distribution and corrected using again a cluster-based procedure.

#### Control analysis

To further ensure the reliability of the t-SPC events identified by our cluster-based permutation analysis, we conducted two control analyses. Firstly, we required that a t-SPC event contain at least two cycles of oscillation at the centroid frequency to be classified as reliable, thus ruling out brief and transitory noise-driven clusters. Secondly, we recognized that surrogate SPC maps generated during the permutation procedure may contain surrogate t-SPC events due to natural cluster tendency arising in small samples of random distribution, which can be mistakenly identified as non-random. To this end, we z-scored the surrogate PPC maps and conducted the same cluster-based permutation, defining surrogate t-SPC events as those that met the same criteria as the original t-SPC events. We then compared the number of observed t-SPC events to that of the surrogate t-SPC events. We also carried out several control analyses to rule out confounding factors that might have influenced the SPC changes we observed: difference in firing rates, differences in ECoG power and a phase reset around speech production onset. Although the SPC pipeline is designed to remove any firing rate bias in the SPC estimation, we sought to investigate genuine firing rate effects by plotting firing rate changes against SPC changes across STN neurons. To ensure that phase estimates were not based on unreliable low amplitude oscillation (during beta suppression), we repeated the analysis and discarded instantaneous phase samples in which the instantaneous power fell below the 10^th^ percentile. We also checked whether bouts of oscillatory power (during theta and gamma increase) biased the SPC estimation by discarding instantaneous phase samples in which the instantaneous power rose above the 90^th^ percentile. We examined the impact of phase resetting of brain oscillations, which can generate event-related activity. To this end, we run two complementary analyses. First, we aligned all the trials to the speech production onset, averaged the ECoG signals across trials to obtain evoked activity, and subtracted this component from individual trials before conducting the SPC analysis. Second, we quantified whether auditory cue or speech production onset reset the phase of the ECoG oscillations. We estimated the event-locked SPC the same way as the SPC, except that ECoG segments were aligned at the auditory cue and speech production onset. Each trial thus contributed one spike to the SPC computation.

#### Statistical analysis

We used the RainCloud library for the visualization of data distributions^76^. Kolmogorov-Smirnov test revealed that the normality assumption of the distribution was rarely satisfied. For this reason, we decided to apply a series of permutation tests (500 permutations) throughout the manuscript whenever definition of a null distribution was methodologically justified. An exception is represented by circular data (e.g. phases) that required the usage of the CircStat toolbox^71^. All results were assessed at statistical significance of *α* = 0.05.

## Lead Contact

Further information and requests for resources should be directed to the Lead Contact, Robert Mark Richardson.

## Data and Code Availability

The data of this study is hosted in the Data Archive BRAIN Initiative (DABI, https://dabi.loni.usc.edu/dsi/1U01NS098969) and is available upon request.

## Acknowledgments

We would like to thank research participants for their generous contribution of time and effort in the operating room and additional experimenters who acquired and organized the data. This work was funded by the National Institute of Health (BRAIN Initiative), through grants U01NS098969, U01NS117836 and R01NS110424 to R.M.R.

## Author contributions

A.B. and W.J.L. wrote experimental code and performed experiments and recorded data. A.B., W.J.L., L.L.H., J.A.F. and R.M.R. designed experiment. R.M.R performed the surgery and supervised the project. P.F. helped to implement the SPC pipeline and contributed to the interpretation of the SPC results. C.N. wrote parts of the discussion and helped to create the 3D visualization of the SPC topography. L.B. contributed to the discussion and interpretation of the results. M.V. analyzed data, prepared figures, and wrote the first draft of the manuscript. All authors discussed results at all stages of the project and revised the manuscript.

