## Supplementary File for "Spike-phase coupling of subthalamic neurons to posterior opercular cortex predicts speech sound accuracy"

#### List of Supplementary Tables

**Table S1: Clinical and demographic patient information.** Summary of the demographic description of the patient cohort, including symptom severity, recordings availability and behavioral performance. M/F: male/female, UPDRS: Unified Parkinson's Disease Rating Scale, NR: value was not recorded in the medical record.

| ID | Age | Sex | Surgery side | UPDRS III OFF | UPDRS III Speech OFF | UPDRS III ON | UPDRS III Speech ON | #task runs | #ECoG strips | MER tracks | #Units | Speech duration, s | # trials | Phonetic accuracy, % |
|---|---|---|---|---|---|---|---|---|---|---|---|---|---|---|
| DBS3001 | 69 | M | Left | 25 | 0 | 10 | 0 | 4 | 1 | MCP | 22 (19) | 1.29 | 344 | 86.61 |
| DBS3002 | 60 | M | Left | 29 | 1 | 16 | 1 | 2 | 1 | MCP | 5 (4) | 0.99 | 360 | 60.34 |
| DBS3003 | 52 | M | Left | 17 | 1 | 3 | 0 | 3 | 2 | MCP | 13 (13) | 1.39 | 480 | 77.60 |
| DBS3004 | 62 | M | Left | 61 | 1 | 36 | 2 | 3 | 1 | MCP | 11 (9) | 1.35 | 360 | 55.00 |
| DBS3008 | 71 | M | Left | NR | NR | NR | NR | 2 | 1 | MCP | 11 (10) | 1.54 | 360 | 74.17 |
| DBS3010 | 70 | M | Left | 34 | 1 | 14 | 1 | 3 | 3 | MCP | 14 (13) | 1.16 | 480 | 78.96 |
| DBS3011 | 66 | M | Left | 30 | 1 | 7 | 0 | 3 | 2 | MCP | 8 (8) | 0.89 | 480 | 69.05 |
| DBS3012 | 70 | M | Left | 43 | 2 | 26 | 2 | 4 | 2 | ACP | 13 (11) | 1.19 | 600 | 84.01 |
| DBS3014 | 71 | M | Left | 30 | 0 | 19 | 0 | 3 | 2 | MCP | 15 (!5) | 1.52 | 480 | 50.42 |
| DBS3015 | 75 | M | Left | 42 | 1 | 24 | 1 | 3 | 1 | MCP | 9 (9) | 1.61 | 480 | 87.29 |
| DBS3017 | 77 | M | Left | 34 | 1 | 26 | 1 | 3 | 1 | MCP | 6 (2) | 1.59 | 345 | 48.83 |
| DBS3018 | 62 | F | Left | 21 | 1 | 13 | 0 | 3 | 2 | MCP | 9 (8) | 1.27 | 480 | 88.08 |
| DBS3019 | 71 | M | Left | 40 | NR | 18 | NR | 3 | 2 | MCP | 13 (7) | 1.13 | 360 | 20.55 |
| DBS3020 | 64 | M | Left | 19 | 0 | 15 | 0 | 2 | 2 | MCP | 7 (7) | 1.28 | 360 | 26.76 |
| DBS3022 | 50 | M | Left | 38 | 1 | 20 | 1 | 2 | 2 | MCP | 9 (9) | 1.05 | 360 | 3.89 |
| DBS3023 | 67 | M | Left | 30 | 2 | 7 | 1 | 3 | 2 | MCP | 11 (11) | 1.24 | 360 | 27.94 |
| DBS3024 | 63 | F | Left | 37 | 1 | 27 | 1 | 2 | 2 | MCP | 11 (8) | 1.13 | 360 | 69.05 |
| DBS3026 | 69 | M | Left | 50 | 1 | 24 | 1 | 2 | 2 | MCP | 7 (5) | 1.23 | 148 | 6.40 |
| DBS3027 | 57 | M | Left | 33 | 0 | 30 | 0 | 2 | 2 | MCP | 9 (9) | 1.37 | 360 | 39.17 |
| DBS3028 | 64 | F | Left | 39 | 2 | 29 | 2 | 3 | 2 | MCP | 18 (16) | 1.48 | 480 | 88.57 |
| DBS3029 | 78 | M | Left | NR | NR | NR | NR | 2 | 2 | MCP | 6 (5) | 1.40 | 240 | 52.08 |
| DBS3030 | 62 | F | Left | 43 | 2 | 30 | 2 | 2 | 2 | MCP | 5 (5) | 2.49 | 240 | 56.25 |
| DBS3031 | 70 | M | Left | 39 | 2 | 26 | 2 | 2 | 2 | MCP | 5 (4) | 1.26 | 240 | 22.38 |
| DBS3032 | 56 | M | Left | 47 | 1 | 27 | 1 | 3 | 2 | MCP | 8 (4) | 1.49 | 360 | 80.39 |
| **Mean (SD)** | **65.44 (7.07)** | **20 M/ 4 F** | - | **35.5 (10.18)** | | | | **2.67 (0.62)** | **1.79 (0.49)** | | **24 5 (211)** | **1.35 (0.41)** | **379.88 (99.49)** | **56.41(26.91)** |

**Table S2: Frequency-wise spatial centroid and peak of spatial density of SPC topographies.** Results are reported for the cortex and subthalamic nucleus.

| | | | | | | | |
|---|---|---|---|---|---|---|---|
| **Cortex** | | | | | | | |
| | **Centroid** | | | | **Peak** | | |
| | **X [mm]** | **Y [mm]** | **Z [mm]** | | **X [mm]** | **Y [mm]** | **Z [mm]** |
| σ | -67.51 | -27.91 | 24.86 | | -69.73 | -49.94 | 12.89 |
| α | -62.23 | -13.04 | 30.4 | | -68.86 | -38.46 | 24.12 |
| β | -65.27 | -10.14 | 28.41 | | -71.14 | -13.78 | 29.80 |
| $\gamma_L$ | -62.66 | -5.45 | 26.74 | | -52.67 | 10.63 | 39.76 |
| $\gamma_H$ | -62.05 | -3.92 | 26.94 | | -49.35 | 16.02 | 41.50 |
| **STN** | | | | | | | |
| | **Centroid** | | | | **Peak** | | |
| | **X [mm]** | **Y [mm]** | **Z [mm]** | | **X [mm]** | **Y [mm]** | **Z [mm]** |
| σ | -13.54 | -15.38 | -6.3 | | -13.65 | -15.23 | -5.4 |
| α | -13.23 | -15.26 | -8.07 | | -13.09 | -15.63 | -8.59 |
| β | -13.32 | -14.16 | -7.64 | | -13.8 | -13.08 | -8.51 |
| $\gamma_L$ | -13.12 | -15.11 | -7.70 | | -15.85 | -16.2 | -6.61 |
| $\gamma_H$ | -12.81 | -15.02 | -7.88 | | -12.33 | -14.27 | -11.77 |

#### List of Supplementary Figures

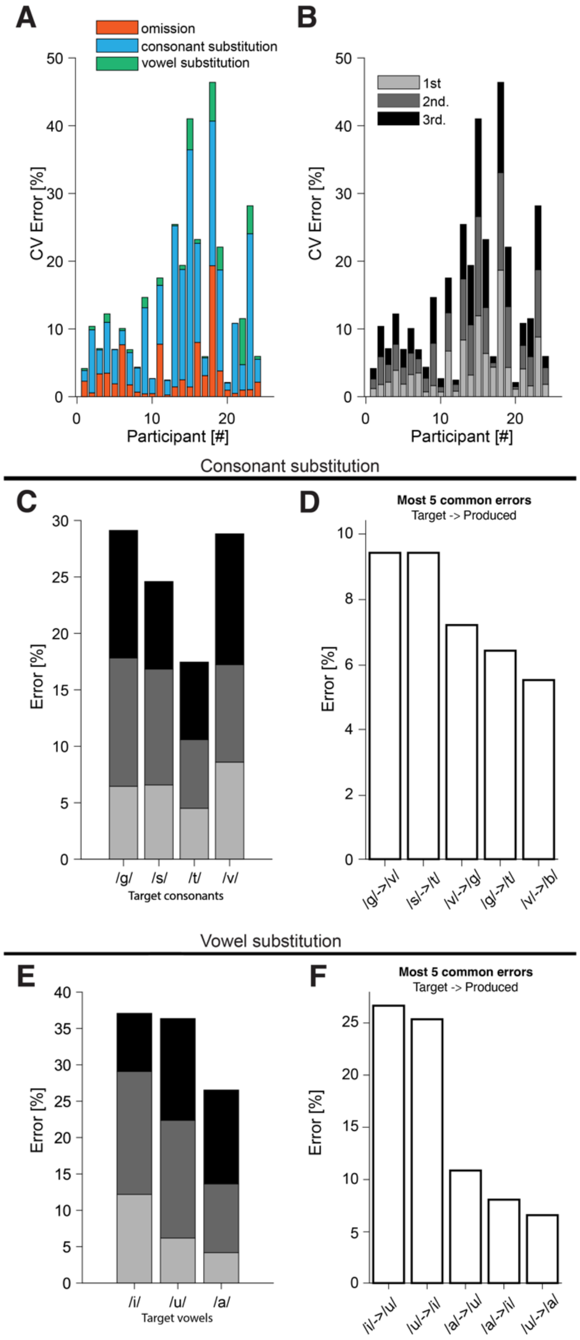


**Figure S1: Characterization of speech errors. (A)** Percentage of speech error types across participants (#errors / # phonemes). Errors were categorized in phoneme omission (orange), consonant substitution (light blue) and vowel substitution (green). **(B)** Proportion of speech errors in the first (light gray), second (gray) and third (black) consonant-vowel (CV) syllable. **(C)** Proportion of consonant substitution error for each target consonant. **(D)** List of 5 common consonant substitution errors across participants. **(E)** Proportion of vowel substitution error for each target vowel. **(F)** List of 5 common vowel substitution errors across participants.

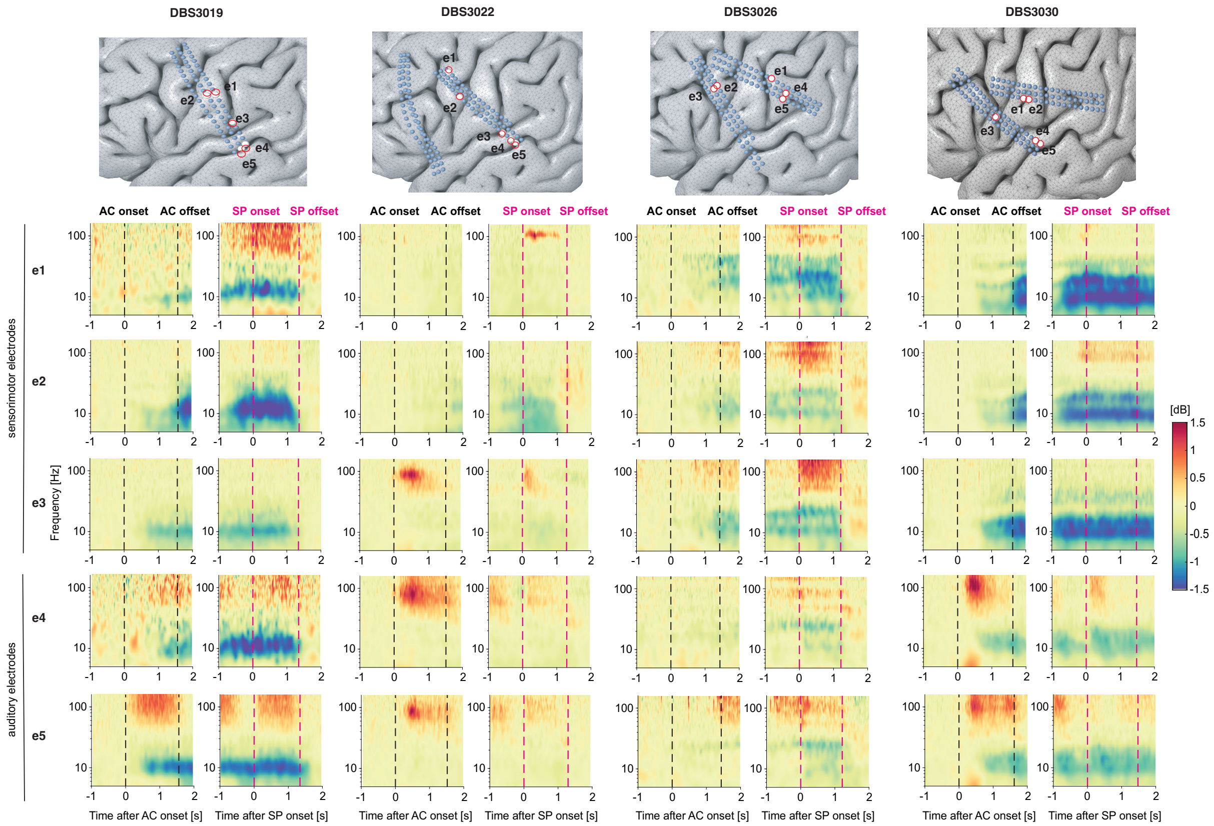


**Figure S2: Electrocorticographic channels from four exemplary participants.** Electrode locations and time-frequency spectrograms locked to auditory cue (AC) (black dashed line) onset and speech production (SP) (magenta dashed line) onset are depicted for each participant. Two representative sensorimotor electrodes (e1, e2) and three auditory electrodes (e3,e4, e5) are shown. Time-frequency spectrograms are normalized with respect to the baseline (during the ITI) and expressed as dB.

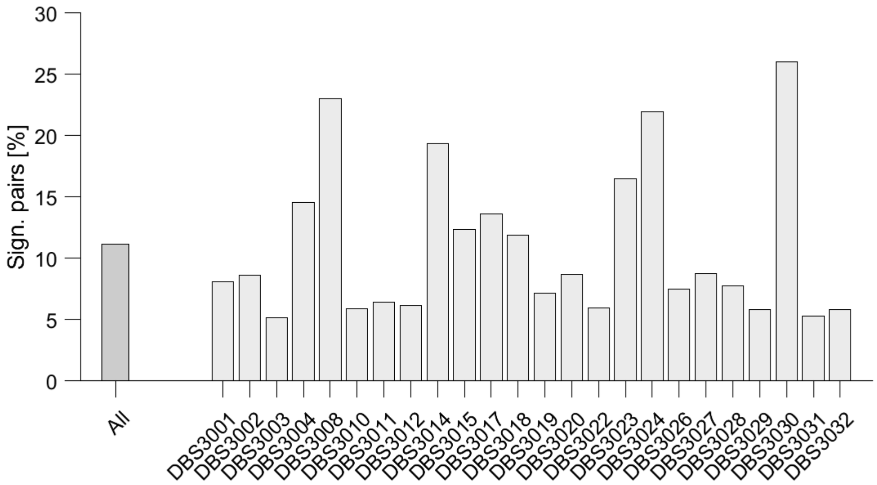


**Figure S3: Percentage of significant SPC pairs across participants.** Results for each participant (light gray) and pooled across all participants (dark gray) are shown.

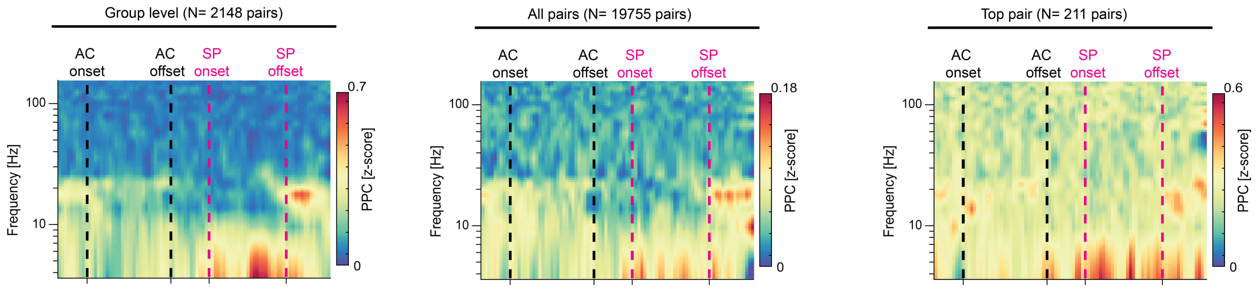


**Figure S4: Comparison of average SPC maps across different subsets of pairs.** Results are consistent whether we averaged only significant SPC maps (N = 2148, same panel of Figure 2A), all pairs (N = 19755) or the most significant SPC map for each unit. Auditory cue (AC) (black dashed line) onset and speech production (magenta dashed line) windows are represented.

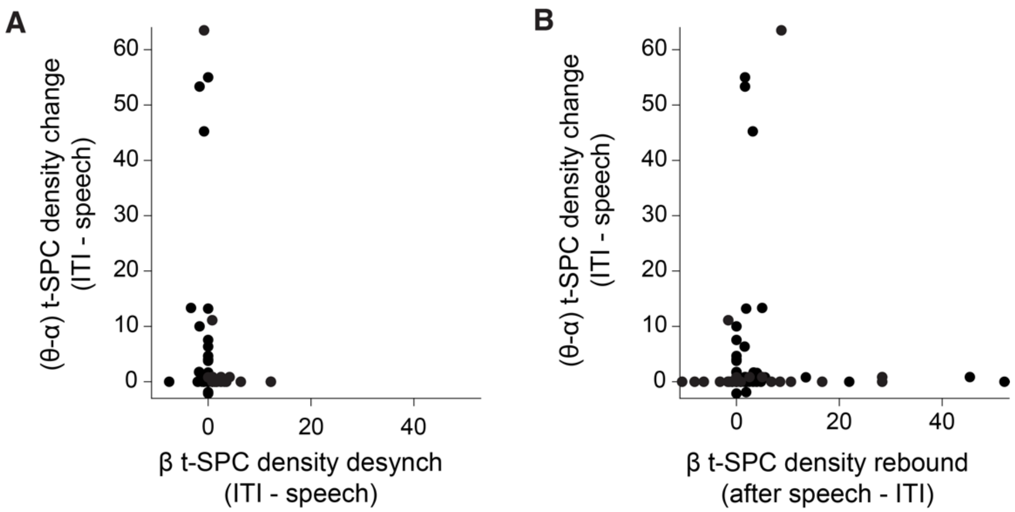


**Figure S5: Neurons are highly specific to only one frequency band.**
Each dot represents the t-SPC spatial density increase in the (θ-α) range plotted against to the decrease in the β-range during the speech production window (**A**), and the increase (rebound) in the β-range after the speech termination (**B**). The observation unit is the neuron (211 dots). The orthogonal relationship highlights the independence between t-SPC density changes in different frequency bands.

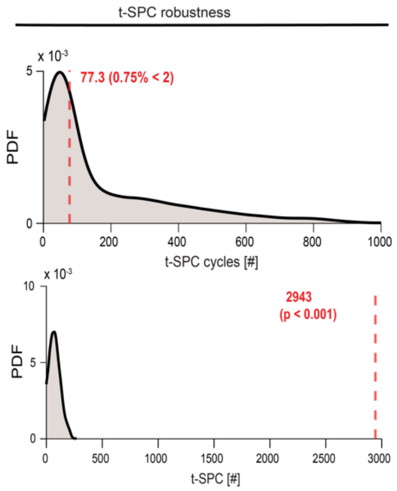


**Figure S6: Robustness analysis of the t-SPC event identification.** **(A)** Distribution of the number of cycles that span the duration of the t-SPC events. Distribution of the number of t-SPC events observed in the permuted SPC maps. Red dashed line depicts the median of the distribution.

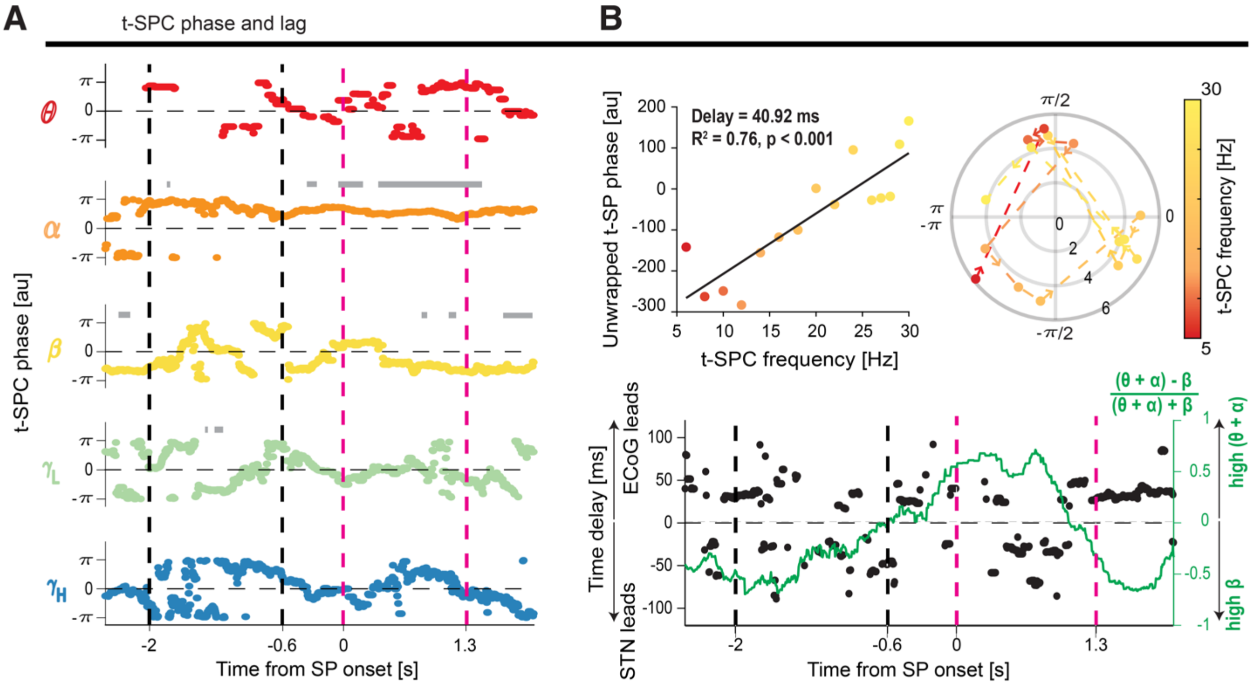


**Figure S7: Phase and time delay relationship (A)** Average preferred phase of firing during the t-SPC events across frequency bands. Gray bar depicts time bins with significant non-uniformity of the preferred phase across all pairs. (**B**) Unwrapped phase values show their progression on a linear scale. The time delay derived from the gradient of the linear fit was 40.92 ms. A permutation test confirmed the significance of the relationship between unwrapped phases and frequency. The sign of the time delays defines the leading activity: positive (ECoG leads STN) and negative (STN leads ECoG). Circular plot shows mean angle (position on circle) and strength of SPC (PPC) of pairs that are significantly locked at frequencies from 5 to 30 Hz (as indicated by the color scale). Time-resolved estimation of the time delay superimposed with the relative occurrence of (θ-α) and β t-SPC events. The sign of this quantity determines the leading frequency band: positive ((θ-α) dominates) and negative (β dominated).

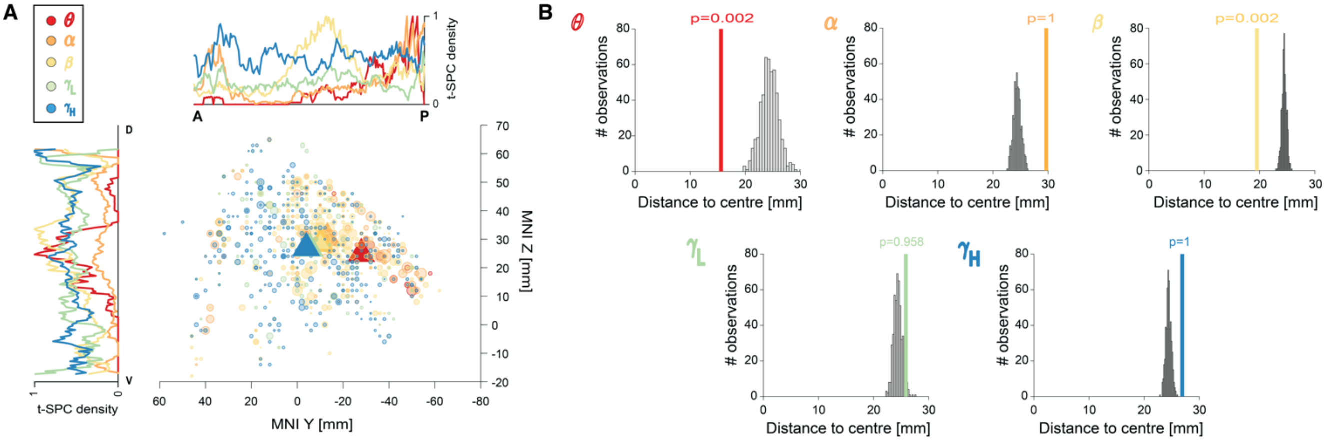


**Figure S8: Spatial aggregation of spectral SPC cortical topographies.** (**A**) Localization of the spike-phase coupling topography of all frequency bands in the sagittal plane (MNI Z vs Y coordinates). Inset plots show the average spatial density mapped along the dorso (D) – ventral (V) and anterio (A) – posterior (P) axes. (**B**) Results of the spatial aggregation analysis to test the focality of the distribution of the SPC topographies. Histogram of the permutation distances and the observed distance (vertical line) are illustrated. Triangles depict the spatial centroid of each topography. θ (red), α (dark orange), β (light orange), $\gamma_L$ (green) and $\gamma_H$ (blue).

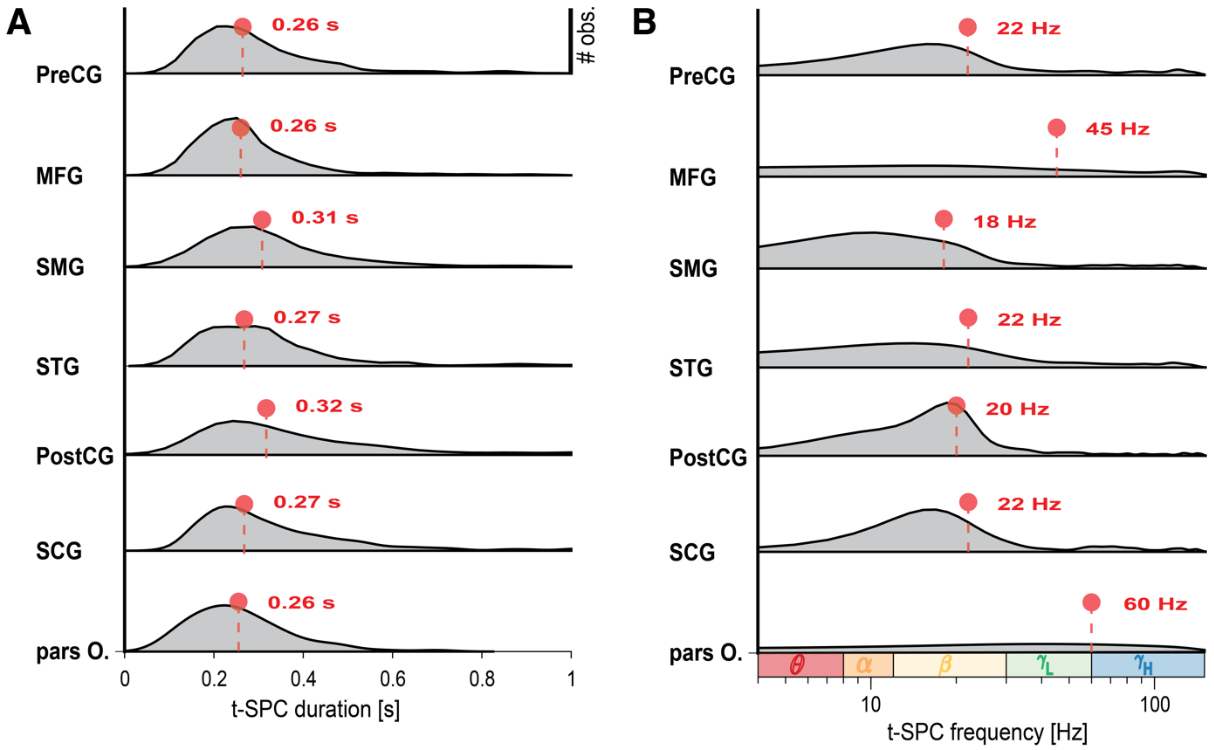


**Figure S9: Cortical heterogeneity of t-SPC events duration and frequency.** Distribution of the t-SPC duration (**A**) and t-SPC frequency (**B**) in seven cortical regions of interest. To augment the readability of the t-SPC frequency distribution, we adopted the logarithmic scale. Dot and dashed lines depict the median of the distribution. $\theta$ (red), $\alpha$ (dark orange), $\beta$ (light orange), $\gamma_L$ (green) and $\gamma_H$ (blue). List of cortical regions of interests: Precentral gyrus (PreCG), Postcentral gyrus (PostCG), Supramarginal gyrus (SMG), Superior temporal gyrus (STG), Middle frontal gyrus (MFG), and the orbital part of the inferior frontal gyrus (pars O.)

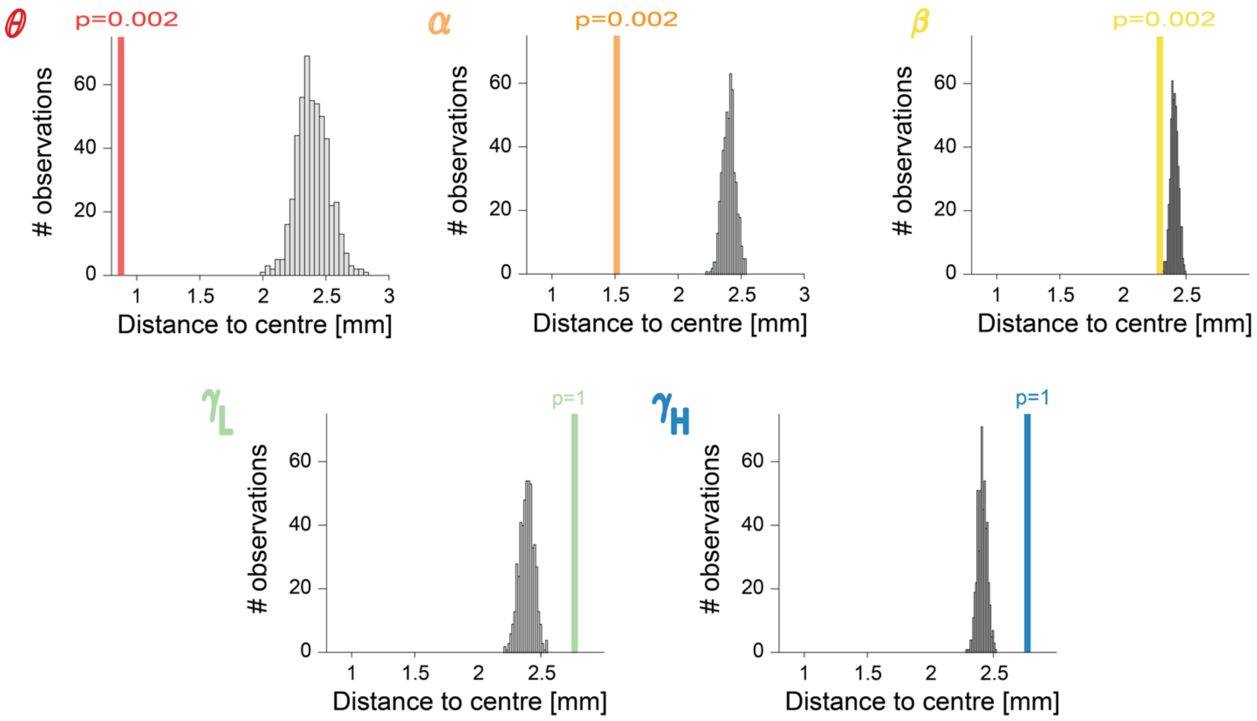


**Figure S10: Spatial aggregation of spectral SPC STN topographies.** Results of the spatial aggregation analysis to test the focality of the distribution of the SPC topographies. Histogram of the permutation distances and the observed distance (vertical line) are illustrated. Triangles depict the spatial centroid of each topography. $\theta$ (red), $\alpha$ (dark orange), $\beta$ (light orange), $\gamma_L$ (green) and $\gamma_H$ (blue).

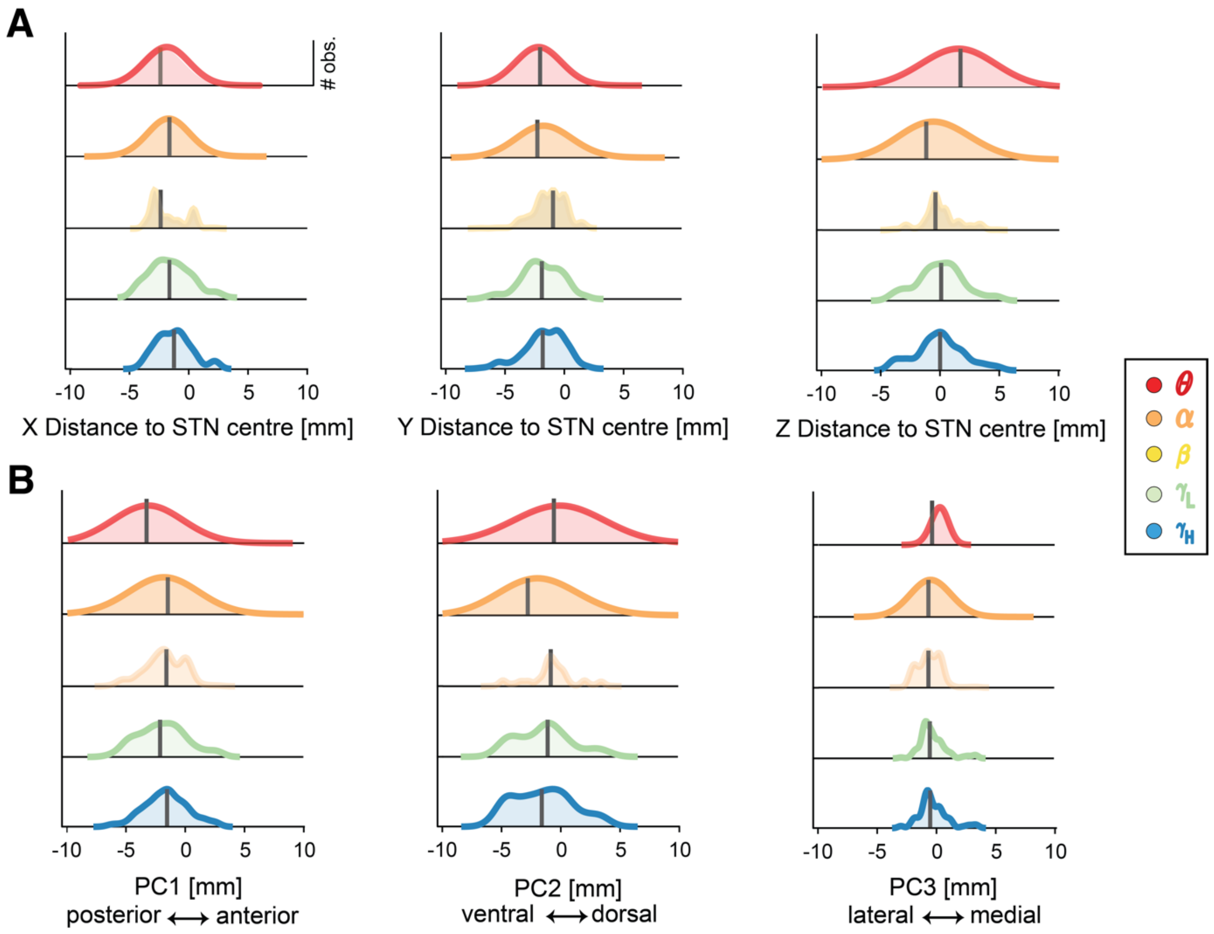

A
### obs.
-10
-5
0
5
10
X Distance to STN centre [mm]
Y Distance to STN centre [mm]
Z Distance to STN centre [mm]
B
PC1 [mm]
posterior ⟷ anterior
PC2 [mm]
ventral ⟷ dorsal
PC3 [mm]
lateral ⟷ medial
θ
α
β
γL
γH

**Figure S11: Localization of the t-SPC events in the STN.** Distribution of the location of the t-SPC events in MNI coordinates (**A**) and principal component axes (**B**) across all frequency bands. Note that principal component scores represent actual physical distances in mm. Black line depicts the median of the distribution. θ (red), α (dark orange), β (yellow), $\gamma_L$ (green) and $\gamma_H$ (blue).

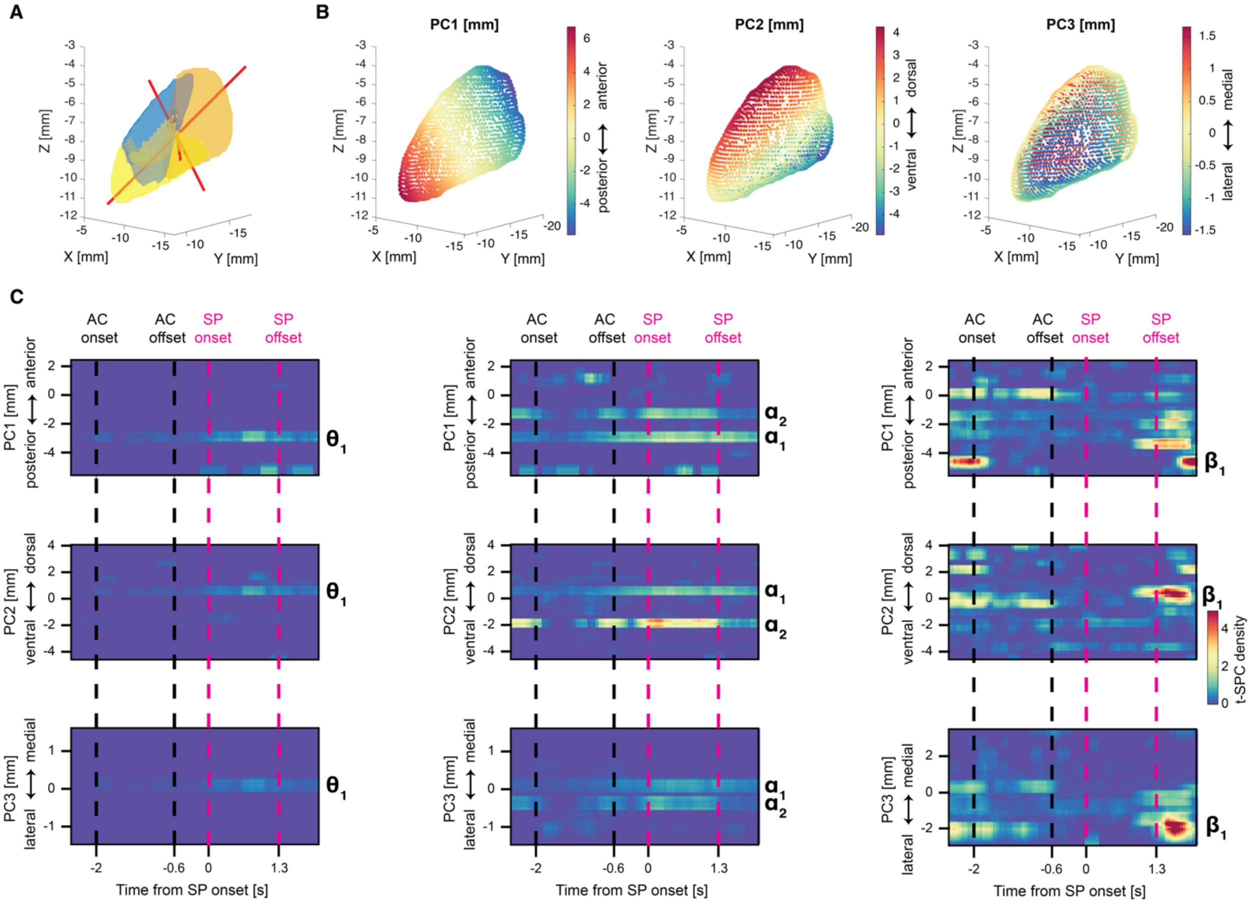

**Figure S12: A new reference of frame to represent STN coordinates.** **(A)** Illustration of the Subthalamic nucleus and its territorial subdivision (motor: orange, associative: yellow and limbic: blue), as depicted by the DISTAL atlas[2]. Direction of the principal component axes is shown as red line. Length of the line is proportional to the explained variance. (**B**) Each principal component has a specific anatomical interpretation: (PC1: anterior-posterior axis, PC2: dorso-ventral axis and PC3: medio-lateral). Note that principal component scores represent actual physical distances in mm. (**C**) Spatial density of the t-SPC events mapped along the three principal component axes. The anatomical reference of frame shows the relative orientation between the dorsal (D), lateral (L) and posterior (P) directions and the first three principal components directions (PC1: anterior-posterior axis, PC2: dorso-ventral axis and PC3: medio-lateral). Black and purple dashed lines denote auditory cue and speech production windows. Cross indicates the spatial centroid of the t-SPC event locations. $\theta_1$, $\alpha_1$, $\alpha_2$ and $\beta_1$ depict the location of peaks of the t-SPC spatial density. Note that principal component scores represent actual physical distances in mm.

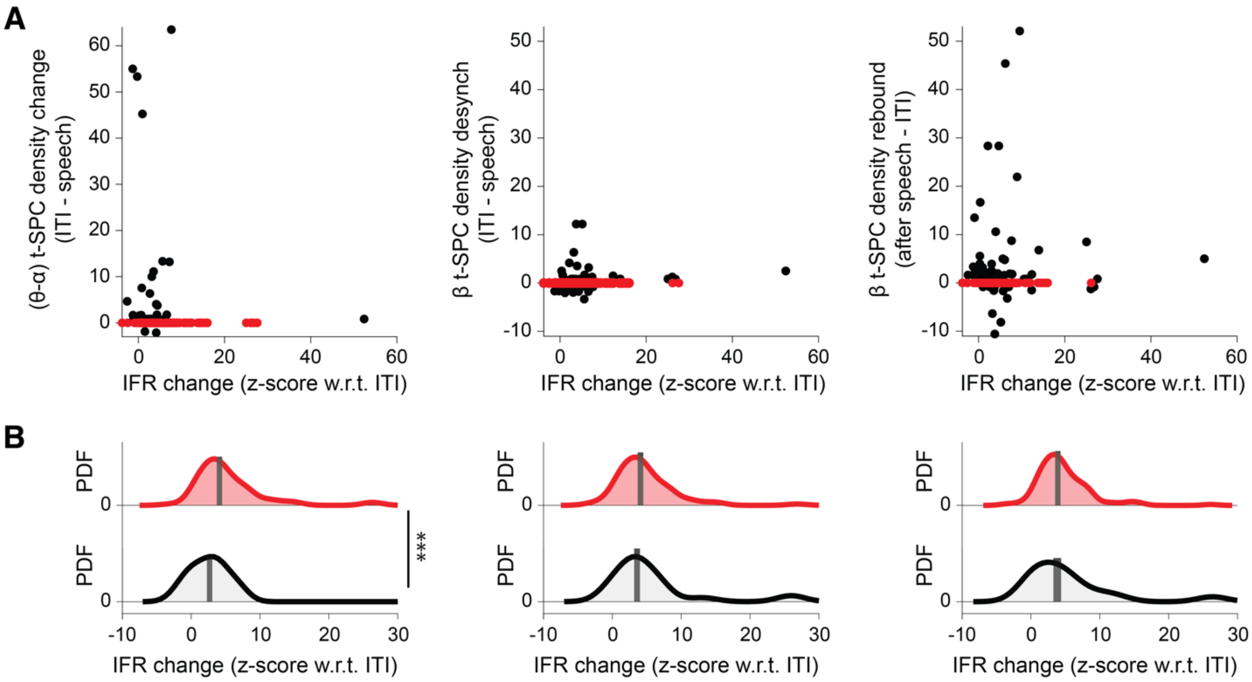


**Figure S13: Relationship between IFR modulation and SPC changes during speech production.** (**A**) Each dot represents the t-SPC spatial density increase in the ($\theta$-$\alpha$) range and decrease in the $\beta$-range during the speech production window, and the increase (rebound) in the $\beta$-range after the speech termination plotted against to the z-score (peak change) of the instantaneous firing rate with respect to the ITI. The observation unit is the neuron (211 dots). Red dots highlight neurons with no change in the t-SPC spatial density. (**B**) Distribution of the z-score change of the instantaneous firing rate in neurons with (black) and without (red) changes in the t-SPC spatial density. Asterisks indicate the significance (***, $p_{perm} < 0.001$) of the permutation test that assesses significant difference between the two distributions. Black line depicts the median of the distribution.

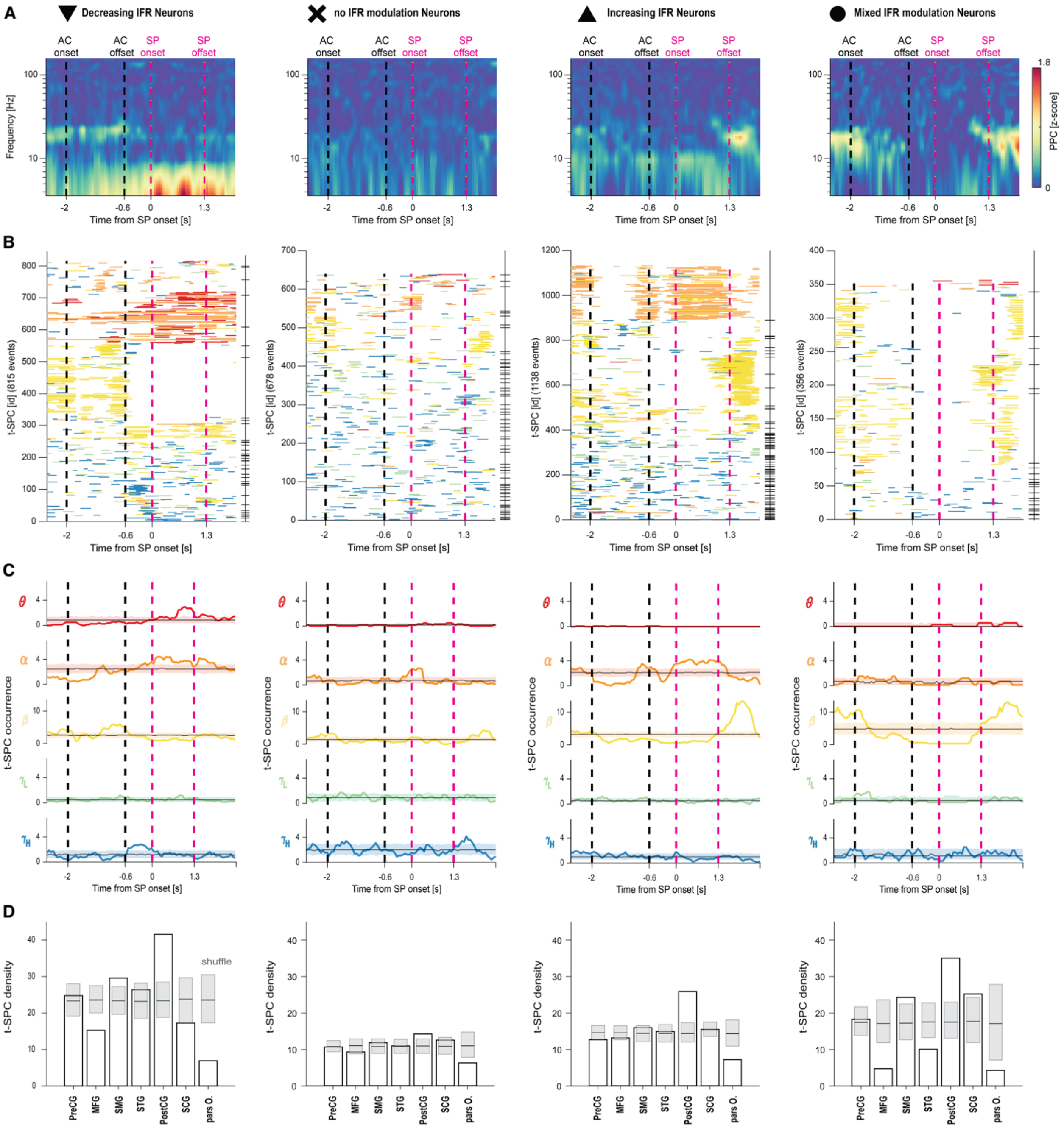
A
Decreasing IFR Neurons
no IFR modulation Neurons
Increasing IFR Neurons
Mixed IFR modulation Neurons
AC onset
AC offset
SP onset
SP offset
Frequency [Hz]
PPC [z-score]
Time from SP onset [s]
B
t-SPC [id] (815 events)
t-SPC [id] (678 events)
t-SPC [id] (1138 events)
t-SPC [id] (356 events)
C
t-SPC occurrence
D
t-SPC density
shuffle
PreCG
MFG
SMG
STG
PostCG
SCG
pars O.

**Figure S14: Spike-phase coupling changes in different instantaneous firing rate categories of neurons.** (**A**) Average of the spike-phase coupling (SPC) maps for significant spike-phase coupling pairs (N = 2148). The pairwise-phase consistency (PPC) index is compared to the permutation distribution and expressed as z-score. (**B**) List of the t-SPC events (N = 2987) sorted by increasing frequency centroid. Horizontal ticks indicate different neurons. (**C**) t-SPC event occurrence grouped by frequency band. Shaded areas illustrate the 5th and 95th percentiles of the permutation distribution for the aggregation test. (**D**) Overall spatial density of the t-SPC events in seven regions of interest, as derived from the Destrieux atlas[26]. Dark gray boxes indicate the 5th and 95th percentile of the permutation distribution for the spatial preference test. Regions with spatial density higher or lower than the permutation distribution are labelled as high or low spatial preference. Black and purple dashed lines denote auditory cue and speech production windows. θ (red), α (dark orange), β (yellow), $\gamma_L$ (green) and $\gamma_H$ (blue).

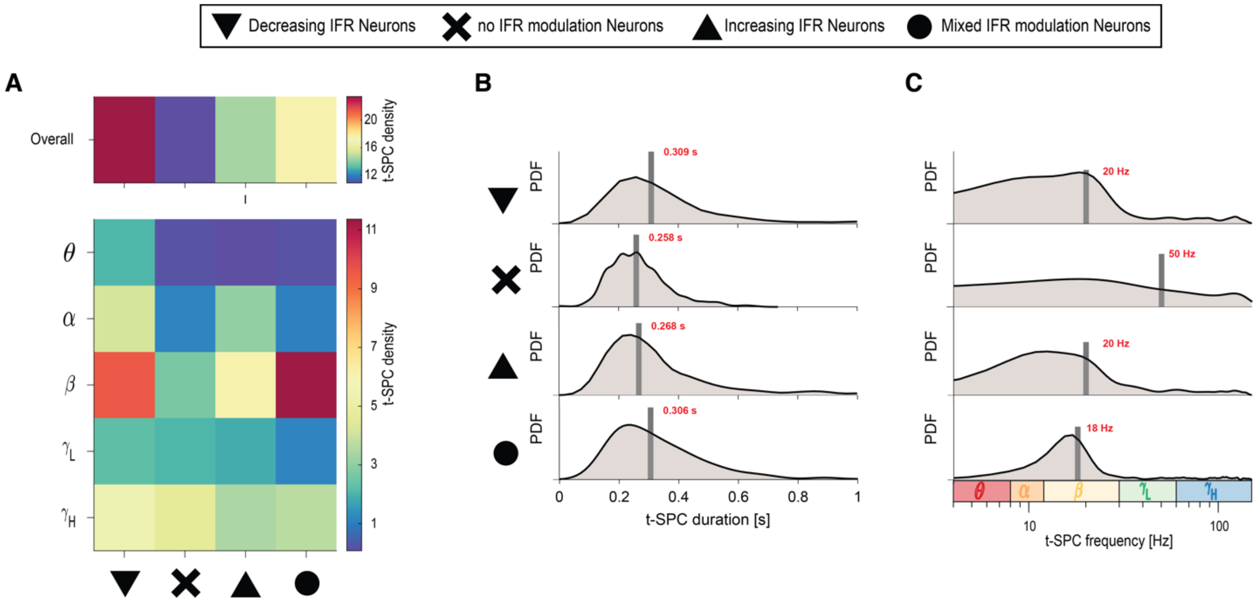


**Figure S15: t-SPC event characteristics across four categories of neurons based on the IFR modulation during the syllable triplet repetition task.** (**A**) Heatmap that illustrates the t-SPC density across different frequency bands θ, α, β, $\gamma_L$ and $\gamma_H$ in each IFR category. Distribution of the t-SPC duration (**B**) and t-SPC frequency (**C**) in each IFR category. To augment the readability of the t-SPC frequency distribution, we adopted the logarithmic scale. List of IFR categories: Decreasing IFR neurons (downward triangle), no IFR modulation neurons (cross), Increasing IFR neurons (upward triangle) and Mixed IFR modulation neurons (circle). Black line depicts the median of the distribution.

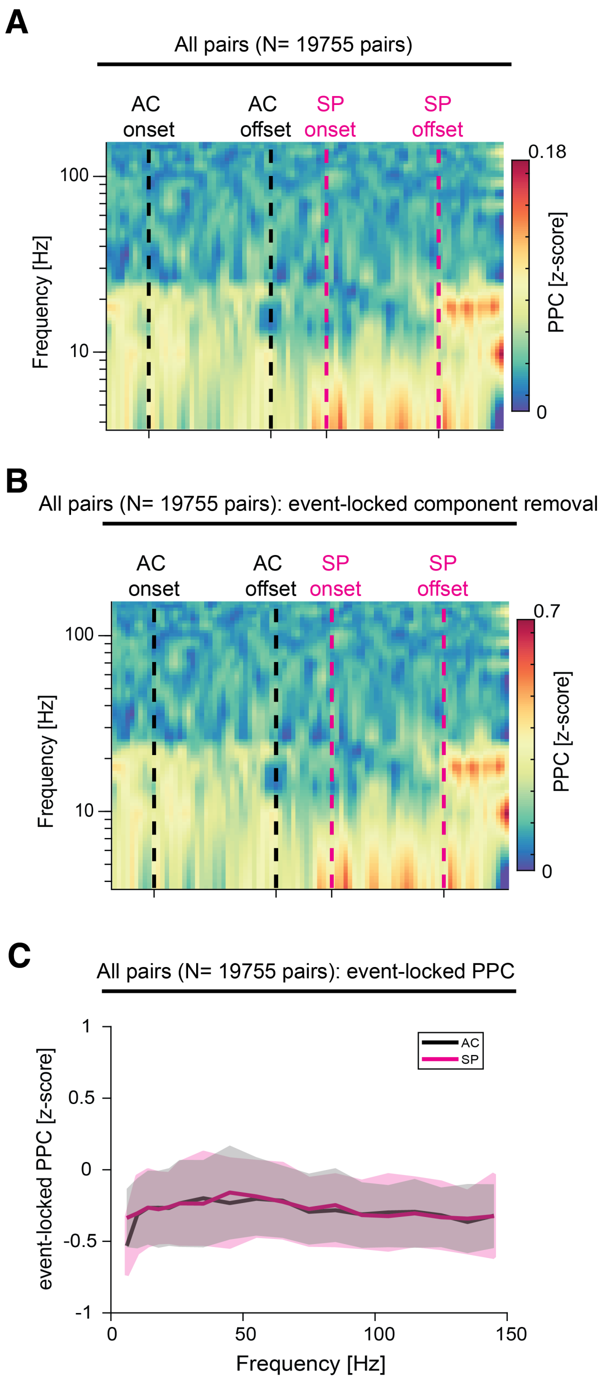


**Figure S16: Event-locked component does not influence the spike-phase coupling estimation.** Results are consistent whether we averaged all SPC maps (N = 19755, same panel of Figure S3B) before (**A**) and after (**B**) the removal of the event-locked component. Auditory cue (AC) (black dashed line) onset and speech production (magenta dashed line) windows are represented. (**C**) Quantification of phase-reset using the same method as used for assessing SPC. We computed event locked SPC by considering as spikes the onset of the auditory cue (black) and speech production (red). Shaded area shows the 95% confidence interval across all pairs (N = 19755).

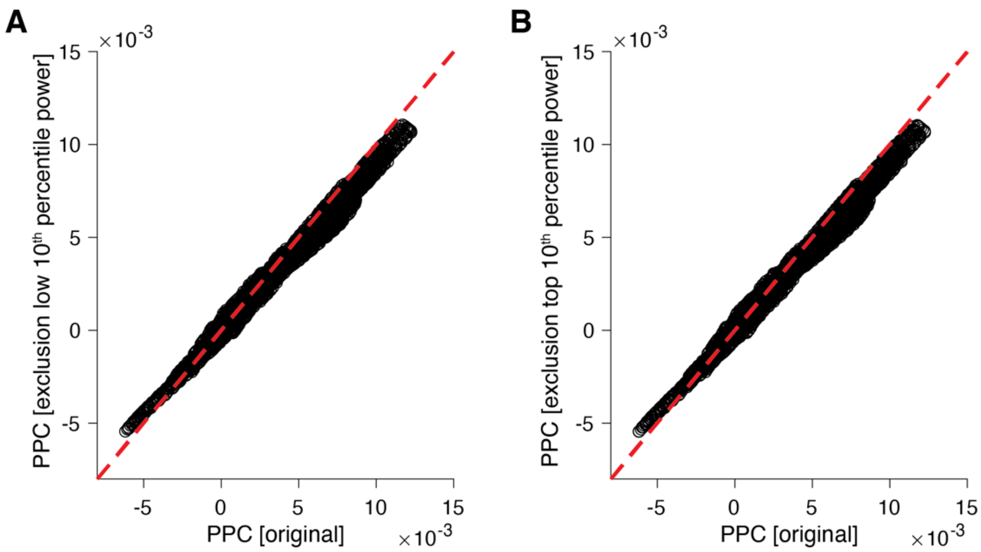


**Figure S17: Calculation of the PPC metric is not affected by extremely high or lower power episodes.** We repeated the calculation of the PPC metric (Methods, SPC pipeline) before and after the removal of periods with low amplitude oscillations (low $10^{th}$ percentile) (**A**) and high amplitude (high $10^{th}$ percentile) (**B**). The red dashed line depicts the identity line.
